# Highly divergent white-tailed deer SARS-CoV-2 with potential deer-to-human transmission

**DOI:** 10.1101/2022.02.22.481551

**Authors:** Bradley Pickering, Oliver Lung, Finlay Maguire, Peter Kruczkiewicz, Jonathon D. Kotwa, Tore Buchanan, Marianne Gagnier, Jennifer L. Guthrie, Claire M. Jardine, Alex Marchand-Austin, Ariane Massé, Heather McClinchey, Kuganya Nirmalarajah, Patryk Aftanas, Juliette Blais-Savoie, Hsien-Yao Chee, Emily Chien, Winfield Yim, Andra Banete, Bryan D. Griffin, Lily Yip, Melissa Goolia, Matthew Suderman, Mathieu Pinette, Greg Smith, Daniel Sullivan, Josip Rudar, Elizabeth Adey, Michelle Nebroski, Guillaume Goyette, Andrés Finzi, Geneviève Laroche, Ardeshir Ariana, Brett Vahkal, Marceline Côté, Allison J. McGeer, Larissa Nituch, Samira Mubareka, Jeff Bowman

**Affiliations:** National Centre for Foreign Animal Disease, Canadian Food Inspection Agency, Winnipeg, Manitoba, Canada; Faculty of Computer Science, Dalhousie University, Nova Scotia, Canada; Sunnybrook Research Institute, Toronto, Ontario, Canada; Wildlife Research and Monitoring Section, Ontario Ministry of Northern Development, Mines, Natural Resources and Forestry, Peterborough, Ontario, Canada; Ministère des Forêts, de la Faune et des Parcs, Québec City, Québec, Canada; Public Health Ontario, Toronto, Ontario, Canada; Canadian Wildlife Health Cooperative, Ontario-Nunavut, Department of Pathobiology, University of Guelph, Guelph, Ontario, Canada; Public Health, Health Protection and Surveillance Policy and Programs Branch, Ontario Ministry of Health; Department of Veterinary Microbiology and Preventative Medicine, College of Veterinary Medicine, Iowa State University, Ames, Iowa, USA; Department of Medical Microbiology and Infectious Diseases, University of Manitoba, Winnipeg; Department of Biological Sciences, University of Manitoba, Winnipeg, Manitoba, Canada; Department of Community Health & Epidemiology, Faculty of Medicine, Dalhousie University, Nova Scotia, Canada; Shared Hospital Laboratory, Toronto, Ontario, Canada; Centre de Recherche du CHUM, Montréal, Québec, Canada; Département de Microbiologie, Infectiologie et Immunologie, Université de Montréal, Montréal, Québec, Canada; Department of Microbiology & Immunology, Western University, London, Ontario, Canada; Department of Biochemistry, Microbiology and Immunology, University of Ottawa, Ottawa, Ontario, Canada; Ottawa Institute of Systems Biology, University of Ottawa, Ottawa, Ontario, Canada; Centre for Infection, Immunity, and Inflammation, University of Ottawa, Ottawa, Ontario, Canada; Sinai Health System, Toronto, Ontario, Canada; Department of Laboratory Medicine and Pathobiology, University of Toronto, Toronto, Ontario, Canada; Environmental and Life Sciences Graduate Program, Trent University, Ontario, Canada

## Abstract

Wildlife reservoirs of SARS-CoV-2 may enable viral adaptation and spillback from animals to humans. In North America, there is evidence of unsustained spillover of SARS-CoV-2 from humans to white-tailed deer (*Odocoileus virginianus*), but no evidence of transmission from deer to humans. Through a biosurveillance program in Ontario, Canada we identified a new and highly divergent lineage of SARS-CoV-2 in white-tailed deer. This lineage is the most divergent SARS-CoV-2 lineage identified to date, with 76 consensus mutations (including 37 previously associated with non-human animal hosts) and signatures of considerable evolution and transmission within wildlife. Phylogenetic analysis also revealed an epidemiologically linked human case. Together, our findings represent the first clear evidence of sustained evolution of SARS-CoV-2 in white-tailed deer and of deer-to-human transmission.

## Main

High consequence coronaviruses, including severe acute respiratory syndrome coronavirus (SARS-CoV), SARS-CoV-2, and Middle East respiratory syndrome coronavirus (MERS-CoV) have putative animal origins and are likely transmitted to humans either directly from reservoir hosts or through intermediates such as civets or camels ^1–6^. This may be followed by sustained human-to-human transmission with ongoing viral adaptation, as we have seen with SARS-CoV-2 and the emergence of variants of concern (VOCs). Additional viral diversity may be gained through inter-species transmission, as was observed during human-mink-human transmission of SARS-CoV-2 ^7^. The emergence of divergent viral lineages can affect viral immunology, biology and epidemiology, thus altering vaccine efficacy, disease severity, and transmission and impacting individual and population health. For example, the Omicron VOC is a highly divergent lineage with 59 mutations across the genome, including 37 in the spike protein, and ultimately led to further global disruption of healthcare systems and societies ^8^. There are several competing theories behind the origin of this VOC; however, transmission analysis of Omicron mutations and molecular docking experiments suggest it may have originated following SARS-CoV-2 transmission from humans to mice and back (i.e., spillback) ^9^ or another, as of yet unidentified, animal reservoir. In addition, animals may contribute to the spread of known VOCs, as demonstrated by regional introduction of a Delta VOC by companion hamsters, with spillback to humans and forward human-to-human transmission ^10^. Establishment of an animal reservoir of SARS-CoV-2 through viral persistence within a susceptible species may lead to repeated spillback events into the human population, with the risk of sustained human-to-human transmission ^11^.

As of March 2022, SARS-CoV-2 has been shown to infect at least 50 non-human mammalian species through both observational and experimental studies in free-living, captive, domestic, and farmed animals ^12–14^. The high degree of homology of the primary SARS-CoV-2 host cell receptor, human angiotensin converting enzyme 2 (hACE2)among certain species may explain this broad host-range ^15^. Zooanthroponosis has been documented in outbreaks of SARS-CoV-2 among farmed (and escaped or feral) mink (*Neovison vison*) in Europe and North America ^16,17^ and pet hamsters (*Mesocricetus auratus*) ^10^. The Netherlands experienced outbreaks of SARS-CoV-2 in mink farms ^18^, and whole genome sequencing (WGS) provided evidence for the emergence of a “cluster 5” variant among farmed mink with a unique combination of mutations, and spillover from mink to humans ^18^. These mutations raised concerns about vaccine efficacy, contributing to the decision in Denmark to depopulate mink ^19^. Similarly, the finding of SARS-CoV-2 in pet hamsters led Hong Kong authorities to cull thousands of animals ^20^.

There have been suggestions (based on experimental data for VOCs) that SARS-CoV-2 host cell receptor tropism has expanded over time, increasing concerns about the potential for spillover into animals. For example, the Alpha variant is capable of infecting mice (*Mus musculus*), and the Omicron variant spike glycoprotein has been shown to bind avian ACE2 receptors, whereas ancestral SARS-CoV-2 did not ^21,22^. This underscores the potential for ongoing expansion of susceptible host species as VOCs continue to emerge.

The white-tailed deer (*Odocoileus virginianus*) is a common and widespread North American ungulate that is susceptible to SARS-CoV-2. An experimental study first showed that deer developed subclinical infection ^23^ A subsequent study found that 40% of free-ranging deer sampled in Michigan, Illinois, New York, and Pennsylvania, USA were positive for SARS-CoV-2 antibodies ^24^Sylvatic transmission among deer and multiple spillovers from humans to deer have also been confirmed ^25–27^. To date, the majority of SARS-CoV-2 in deer has been similar to prevalent lineages circulating among humans in the same region, suggesting multiple, recent spillover events ^26,27^. However, a divergent Alpha VOC has been reported in deer, providing some evidence of adaptation to a new host species ^28^. There was no previous evidence of deer-to-human transmission of SARS-CoV-2.

In response to evidence of free-ranging white-tailed deer (WTD) infection with SARS-CoV-2, the potential establishment of a deer reservoir, and the risk of deer-to-human transmission, we initiated a SARS-CoV-2 surveillance program of WTD in Ontario, Canada to better understand SARS-CoV-2 prevalence in regional WTD. This work provides evidence of extensive parallel evolution of SARS-CoV-2 in a deer population in Southwestern Ontario with unsustained deer-to-human transmission.

## Results

### Highly divergent SARS-CoV-2 found in deer

From 1 November to 31 December 2021, 300 WTD were sampled from Southwestern (N=249, 83%) and Eastern (N=51, 17%) Ontario, Canada during the annual hunting season (Fig. 1). The majority of sampled WTD were adults (94%) with comparable numbers of females (N=135, 45%) and males (N=165, 55%). We collected 213 nasal swabs and tissue from 294 retropharyngeal lymph nodes (RPLN) which were tested for the presence of SARS-CoV-2 RNA using RT-PCR.

**Fig. 1:**
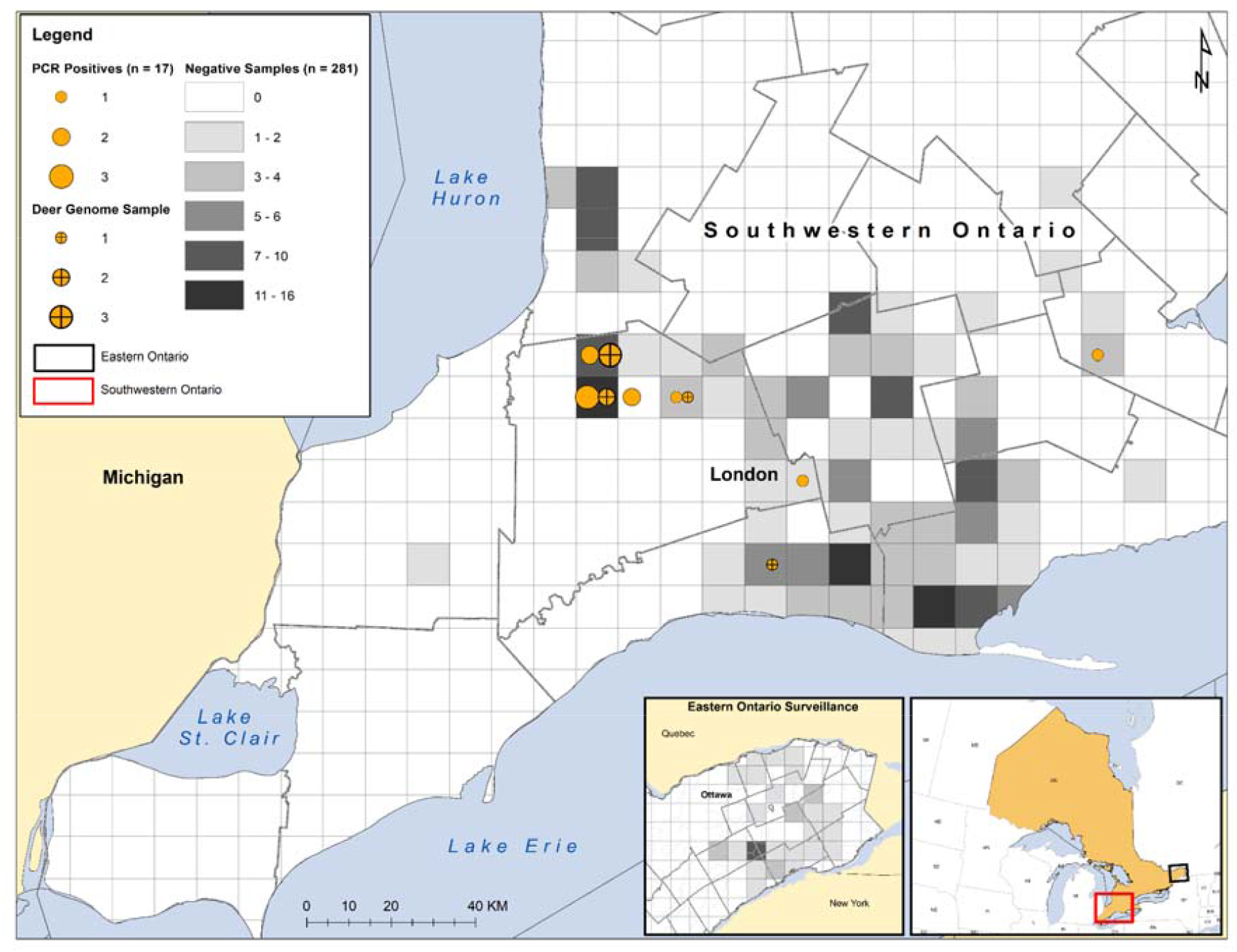
SARS-CoV-2 RNA detection in WTD sampled in Southwestern and Eastern Ontario in 2021. Circle size indicates the relative number of positive WTD (n=17/298), with crosses showing samples from which viral genomes were recovered (n=7). The detailed map depicts Southwestern Ontario (the red rectangle on the inset map). SARS-CoV-2 RNA was not detected in samples from Eastern Ontario.

Five of 213 (2.3%) nasal swabs were positive by two independent RT-PCR analyses at separate institutes (UTR and E gene Ct<40; and E and N2 gene Ct<36). Sixteen RPLN were also confirmed by PCR. Overall, SARS-CoV-2 RNA was detected in 21 samples representing 6% (17/298) of hunter-harvested WTD; all positive animals were adult deer from Southwestern Ontario and the majority (65%) were female (Fig.1, Table S1). Two deer were excluded from further analysis due to indeterminate RPLN results with no corresponding nasal swab.

From the 5 positive nasal swabs, 3 high-quality SARS-CoV-2 consensus genomes were recovered using a standard amplicon-based approach. All samples were also independently extracted and sequenced using a capture-probe-based approach for confirmation. By combining the amplicon and capture-probe sequencing data, 5 high quality genomes (WTD nasal swabs: 4581, 4645, 4649, 4658, and 4662) and 2 partial genomes (WTD RPLNs: 4538, 4534) were recovered (Table S1). The samples were negative for human RNAse P by PCR and the majority (median 66%; Table S1) of non-SARS-CoV-2 reads mapped to the WTD reference genome, demonstrating that contamination from human-derived SARS-CoV-2 sequences was highly unlikely.

Maximum-likelihood and parsimony-based phylogenetic analyses showed these WTD genomes formed a highly divergent clade within the B.1 PANGO lineage/20C Nextstrain clade (100% Ultrafast Bootstrap [UFB]) that shared a most recent common ancestor (MRCA) between December 2020 and September 2021. The B.1 lineage encompasses significant diversity and was the genetic backbone from which the Beta VOC, Epsilon and Iota variants under investigation (VUIs), and significant mink (*Neovison*) outbreaks emerged (Fig. 2). The WTD clade forms a very long branch with 76 conserved nucleotide mutations relative to ancestral SARS-CoV-2 (Wuhan Hu-1) and 49 relative to their closest common ancestor with other genomes in GISAID (as of March 2022). The closest branching genomes in GISAID are human-derived sequences from Michigan, USA, sampled approximately 1 year prior (November/December 2020), which were inferred to share an MRCA with the WTD clade between April and October 2020. These sequences in turn are closely related to a mixed clade of human and mink sequences from Michigan collected in September/October 2020.

**Fig. 2:**
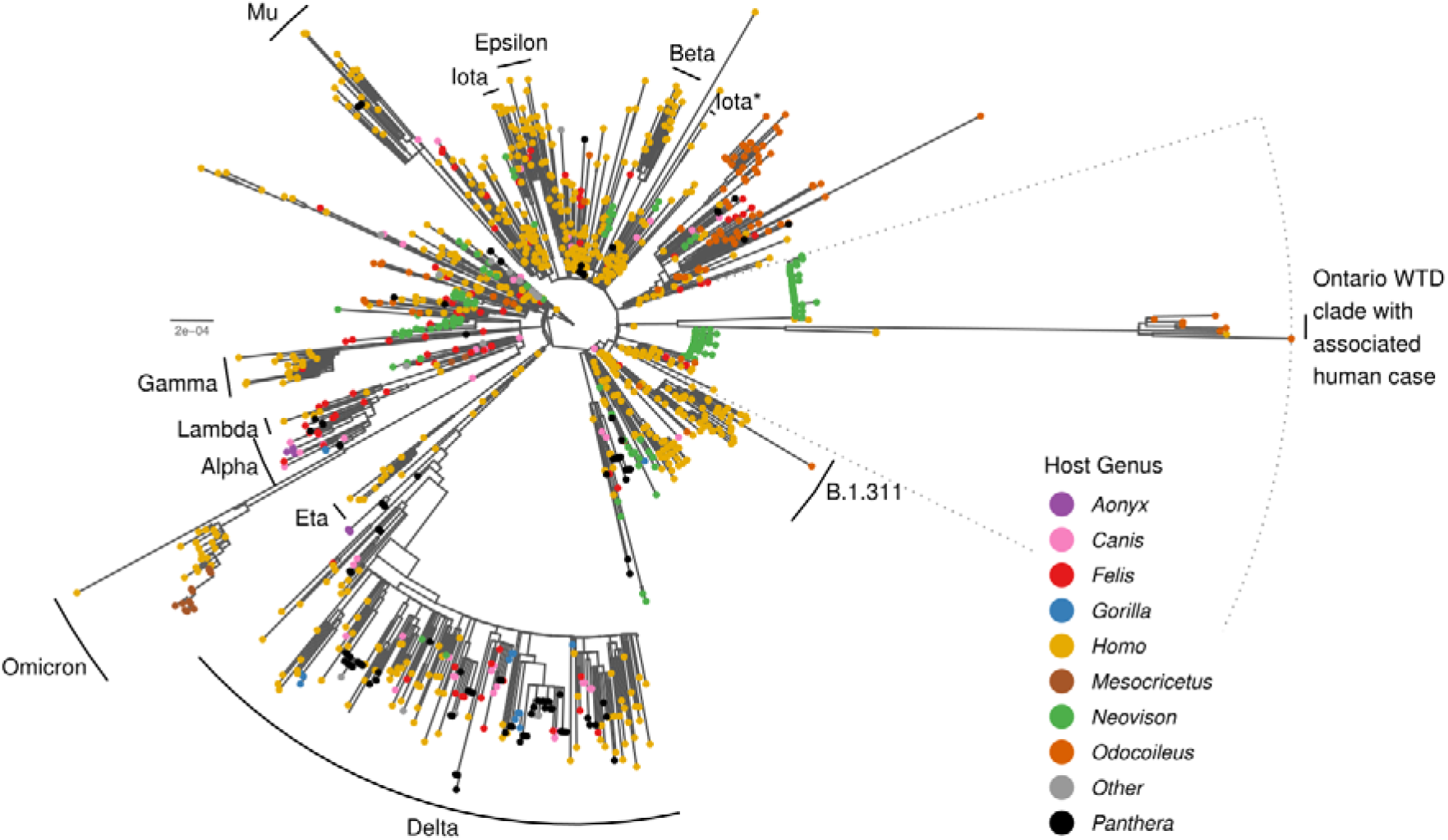
Maximum-likelihood (ML) phylogeny of WTD-derived viral genomes (and associated human sample) and a representative sample of the global diversity of human and animal-derived SARS CoV-2 (n=3,645). This phylogeny under-samples human sequences to maximize representation of animal-associated diversity. VOCs/VUIs within the tree are annotated and nodes are coloured by host genus (as indicated in the legend). The dotted line indicates the samples selected for the local ML analysis (Fig. 3).

Given the distorting effects of long-branch attraction and incomplete sampling, there is a degree of uncertainty in the phylogenetic placement of the WTD samples. However, the geographical proximity (Michigan is adjacent to Southwestern Ontario) and the mix of human and other animal cases (e.g., human and mink cases) provide compelling evidence supporting this placement. Given the degree of divergence and potential for phylogenetic biases, we conducted three analyses to examine the possibility of recombination. Using 3Seq ^29^, bolotie ^30^, and HyPhy’s ^31^ Genetic Algorithm Recombination Detection method ^32^ with datasets representative of human and animal SARS-CoV-2 diversity in GISAID (as of March 2022) there was no indication of recombination within, or having given rise, to this clade.

### Potential deer-to-human transmission

Our phylogenetic analysis also identified a human-derived sequence from Ontario (ON-PHL-21-44225) that was highly similar (80/90 shared mutations; Table S2) and formed a well-supported monophyletic group (100% ultrafast bootstrap support or UFB) with the WTD samples (Fig.3). The small number of samples and relative diversity within the WTD clade make it difficult to determine the exact relationship between the human sample and other WTD samples (78% UFB for a most recent common ancestor with 4658). However, global (Fig. 2) and local (Fig. 3) ML analyses and an Usher-based ^33^ (Fig. S1) parsimony analysis all support this human sample belonging to the WTD clade.

**Fig. 3:**
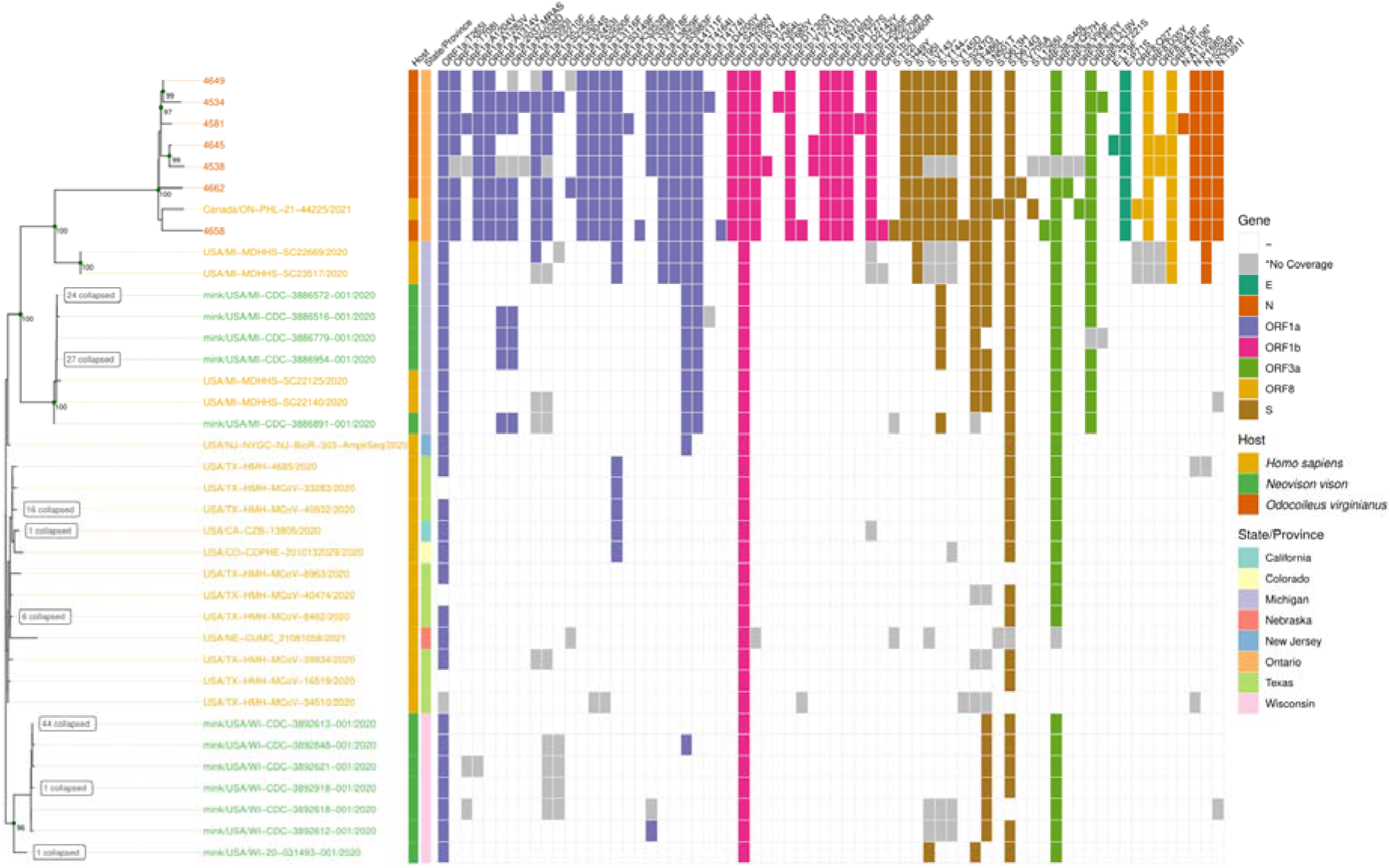
ML phylogeny of 157 genomes selected from global phylogeny annotated with the presence/absence of amino acid mutations relative to SARS-CoV-2 Wuhan Hu-1. Genomes were selected to explore the relationship between Ontario WTD, the related Ontario human sample, and closest B.1 human and mink samples from Michigan, USA (dotted segment in Fig. 2). Internal nodes in the phylogeny are annotated with ultrafast bootstrap support values >=95% and leaves with identical amino acid profiles were collapsed as indicated. Host species for each sample is shown by the leaf label colour and first annotation column as per the legend, with geographic location in the second annotation column. Amino acid mutations are coloured by corresponding gene with grey indicating sites that were too poorly covered to determine presence/absence (e.g., sites in partial WTD genomes 4538, 4534).

The human-derived viral sequence also has a plausible epidemiological link to the WTD samples since it was collected in the same geographical region (Southwestern Ontario), during the same time period (autumn 2021). The human case had known close contact with deer in the week prior to symptom onset and had no known contact with any individuals that had tested positive for SARS-CoV-2 prior to or after contact with deer. At the time of the human case detection, the Ontario COVID-19 Genomic Network aimed to sequence 100% of eligible confirmed PCR positive SARS-CoV-2 samples collected from human cases, and no other genetically related human-derived samples were identified. It should be noted that not all requested human samples are received or successfully sequenced, and the Omicron surge necessitated a reduction in the proportion of human-derived SARS-CoV-2 sampled for sequencing in Ontario in late 2021 ^34^.

### Zoonosis associated mutations

Using the five high-quality complete WTD sequences and related human-derived sequence, we analyzed the prevalence of mutations across GISAID in general as well as within VOC and animal-derived samples (Table S2). Of the 76 mutations shared among the 5 high quality WTD and associated human sequences, 51 are in ORF1ab (with 11 and 9 each in Nsp3 and Nsp4, respectively) and 9 are in the spike (S) gene. The 6 non-synonymous mutations in S correspond to a 6-nucleotide deletion (V143-Y145), and 5 substitutions (H49Y, T95I, F486L, N501T, D614G). These mutations have been observed in animal-derived viral sequences. Not all S mutations were conserved across the entire Ontario WTD clade; S:613H and S:L1265I were found only in the human sample, 3 other non-synonymous mutations were found in either 4658 (T22I and S247G) or 4662 (V705A) WTD samples, and there was a frameshift in 4662 S:L959.

Many non-synonymous mutations had previously been identified in WTD including 16 in at least 3/5 of the WTD samples, S:613H and ORF8:Q27* (associated human sample only), and S:T22I (1/5 ON WTD samples only, but also noted in Delta-like SARS-CoV-2 from deer in Québec ^3^). However, there were also 5 conserved non-synonymous mutations that had not been previously observed in WTD and were relatively rare in GISAID (<1000 sequences as of 14 March 2022): ORF1a:insertion2038N/MRASD (n=32, including 31 mink from Michigan, USA), ORF1b:V364L (G14557T, n=442, all human sequences), S:F486L (T23020G, n=455), ORF3a:L219V (T26047G, n=886), and ORF10:L37F (C29666T, n=0).

### Mutational signatures of deer adaptation

We identified a potentially elevated mutation rate (3.7×10^−3^ vs 0.9×10^−3^ substitutions per site per year) in the Ontario WTD clade (Fig. 4B) relative to other SARS-CoV-2 based on a root-to-tip regression of the global phylogeny (Fig. 2). To characterize signatures of selection within the Ontario WTD clade relative to background B.1 samples, we generated codon-alignment phylogenies for S, E, M, N, ORF3a, ORF6, and ORF1ab sequences and applied selection analysis methods from HyPhy ^31^. The adaptive Branch-Site Random Effects Likelihood ^35^ and Branch-Site Unrestricted Statistical Test for Episodic Diversification ^36^ branch-site methods identified no WTD clade branches with evidence of episodic diversifying positive selection relative to background B.1 branches. Interestingly, the ORF1ab analysis identified significant relaxation of selection amongst the WTD clade (p=0.0032). These signatures of neutrality were further supported by the even distribution of conserved mutations in proportion to gene/product length (e.g., ORF1ab, 54.1 expected vs 51 actual; and spike, 9.7 expected vs 9 actual) (Extended Data Fig. 1). Together, this suggests sustained viral transmission with minimal immune pressure in a susceptible animal population such as WTD. However, further investigation into the host response and disease course of SARS-CoV-2 in WTD is required to confirm these inferences.

**Fig. 4:**
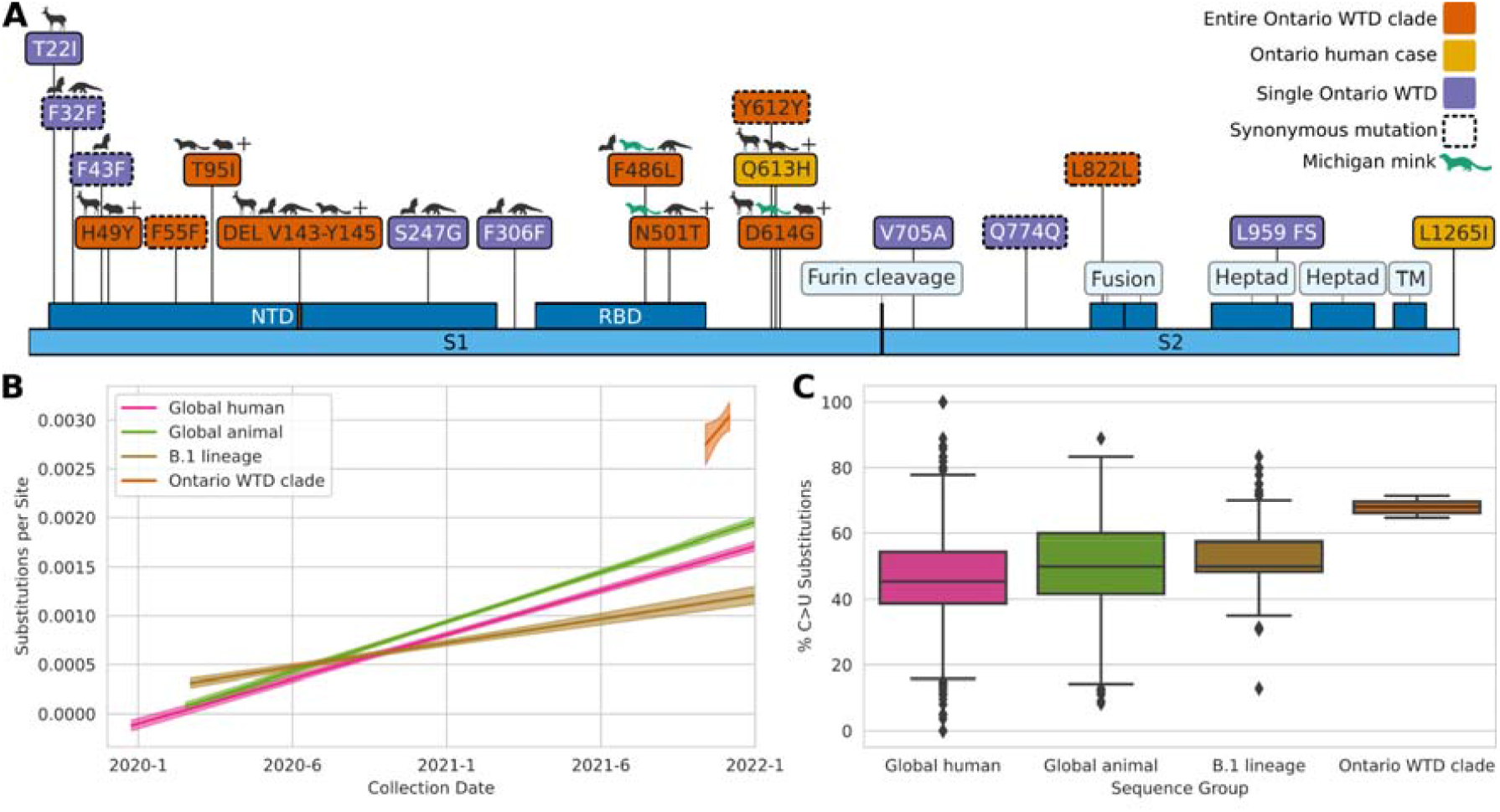
Analysis of the Ontario WTD clade relative to other animal-derived genomes, the ancestral B.1 lineage, and the global SARS-CoV-2 diversity. (A) Amino acid mutations across the spike protein in the Ontario WTD. Amino acid changes present in all 5 Ontario WTD sequences and associated human case (orange); only in the human sample (yellow); and only in a single WTD genome (purple). Animal symbols indicate mutations in bat, deer, pangolin, and hamster-derived SARS-CoV-2 sequences. “+” indicates presence of the mutation in additional animal species, and green indicates those in Michigan mink samples. Spike annotations were derived from UniProt P0DTC2 (DEL: deletion, FS: frameshift, TM: transmembrane, RBD: Receptor binding domain, NTD: N-terminal domain) and are not shown to scale. (B) Root-to-tip regression analysis based on the representative SARS-CoV-2 diversity in the global maximum-likelihood phylogeny (Fig. 2). Substitutions per site per year trends (and 95% confidence intervals) from ordinary least squares regression analyses are shown for all human samples (0.9×10^-3^ to 1.0×10^-3^), animal-derived samples (1.0×10^-3^ to 1.1×10^-3^), the B.1 lineage (0.4×10^-3^ to 0.6×10^-3^), and the Ontario WTD clade (0 to 8×10^-3^). (C) Consensus substitutions (%) corresponding to a change from a reference C allele to an alternative U allele. This was calculated from consensus sequences across a subsample of global human SARS-CoV-2 diversity (earliest and most recent genomes from each PANGO lineage, n=3,127), global animal diversity (all animal genomes currently in GISAID, n=1,522), B.1 lineage (all genomes assigned to this lineage in GISAID as of January 2022, n=206), and the Ontario WTD clade (5 WTD genomes and associated human case).

Changes in the mutational signature of SARS-CoV-2 could be used to trace and understand its spread between human and non-human hosts, and patterns or categories of mutations can provide insights into mechanistic processes (e.g., positive selection, error-prone RNA-dependent RNA polymerase activity or host cell modification through RNA editing). An analysis of base substitution frequencies within the Ontario WTD clade (and associated human sample) showed an elevated proportion of mutations involving C>U changes relative to other global, B.1 lineage, and animal-derived viral sequences (Fig. 4C, Extended Data Fig. 2). Further investigation using a robust non-parametric alternative to MANOVA, dbWMANOVA, found that the mutational spectra between human, deer, and mink hosts differs significantly when considering all clades (W*d = 91.04, p < 0.001) and within Clade 20C, which contains the B.1 lineage and Ontario sequences (W*d = 160.47, p < 0.001) (Table S3, Extended Data Fig. 3). Principal component analysis (PCA) indicated that the majority of this variation (62.2%) corresponded to C>U (PC1) and G>A (PC2) frequencies. Notably, when compared to the recently collected WTD virus samples from Quebec, the location of the Ontario WTD clade (and associated human case) suggests that the Ontario viruses have been evolving within deer (Extended Data Fig. 4).

Analyses of genome composition and codon usage bias may provide information on virus evolution and adaptation to host. We assessed whether the codon usage signatures of the Ontario WTD clade is similar to that of other SARS-CoV-2 sequences (samples isolated from Wuhan-Hu-1, Quebec WTD, mink from Canada and the USA), cervid viruses (Epizootic hemorrhagic disease virus (EHDV), Cervid atadenovirus A, elk circovirus), and the *Odocoileus virginianus* (WTD) genome. No apparent differences were observed in codon usage bias between the Ontario WTD clade and other SARS-CoV-2 sequences across the entire coding region of the viral genome. Although some similarity in codon usage bias to Cervid atadenovirus A was observed, generally there were clear differences between SARS-CoV-2 and non-SARS-CoV-2 sequences (Table S4).

### Virus isolation and S antigenicity

Virus isolation was carried out using Vero E6 cells expressing human transmembrane protease serine 2 (TMPRSS2) with cathepsin L knocked out. At 4 days post-infection (dpi) cytopathic effect of 50% or less of the cell monolayer was observed for four of the samples (4581, 4645, 4658, and 4649) and virus supernatants were harvested. Confirmatory quantitative PCR for SARS-CoV-2 was carried out using 5 ‘UTR and E gene targets. The cycle thresholds for the four isolated samples (1/7 dilutions), 4581, 4645, 4658, and 4649, were 14.89, 16.39, 12.80, 13.89 and 16.17, 24.18, 13.06, 13.91, respectively for 5 ‘UTR and E amplicons respectively. Confirmatory sequencing was carried out successfully for isolates from samples 4581, 4645, and 4658. These showed only minor frequency variations of one SNP change (4581: gain of ORF3a Pro42Leu) or two SNP changes (4658, loss of n.13T>C and gain of ORF1a p.His3125Tyr) compared to the original swab consensus sequences.

Considering that S-gene mutations may lead to immune evasion to antibody responses generated by vaccination or previous infection, we measured spike recognition and neutralizing activity of plasma from vaccinated recipients or convalescent individuals to S glycoproteins identified in this study. Cells were transfected with codon-optimized S expression constructs corresponding to the S genes of samples 4581/4645, 4658, ON-PHL-21-44225, or SARS-CoV-2 S:D614G or Omicron (BA.1) and incubated with sera to analyze antibody recognition of S (Fig. 5A). We found that all WTD S were recognized to similar extent to the S:D614G by sera from vaccinated or convalescent individuals, while Omicron S was less recognized overall (Fig. 5A; Table S5). In addition, lentiviral pseudotypes were incubated with serial dilutions of sera and neutralization half maximal inhibitory serum dilution (ID50) was determined (Fig. 5B; Table S6). We found that sera from vaccinated recipients, either after two or three doses, and from convalescent individuals efficiently neutralized all WTD S proteins, unlike Omicron which required three doses for neutralization (Fig. 5B). Importantly, we did not observe a difference between the ability of sera to neutralize SARS-CoV-2 D614G or any of the Ontario WTD SARS-CoV-2. Taken together, these results suggest that the WTD S gene mutations do not have significant antigenic impact.

**Fig. 5:**
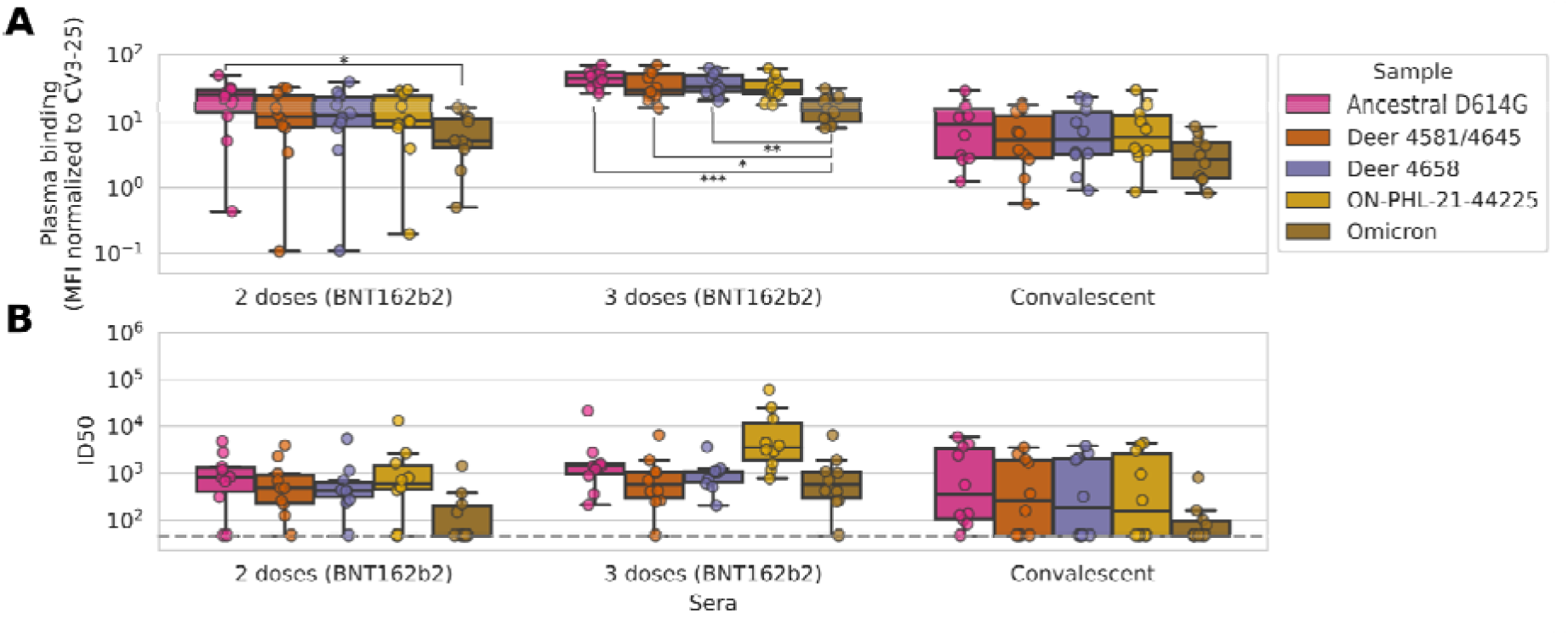
Plasma recognition and neutralization of the Ontario WTD spike glycoproteins and pseudotypes. (A) 293T cells transfected with plasmids encoding the indicated spike variants were incubated with 1:250 diluted plasma from vaccinated (two or three doses of BNT162b2), convalescent, or naïve individuals (n=10 for each group) or with the conformationally-independent anti-S2 CV3-25 antibody, followed by staining with fluorescently labeled anti-human IgG and flow cytometry analysis. Mean fluorescence intensity (MFI) was normalized by surface expression of spike variants based on CV3-25 binding (Table S5). (B) Lentiviral pseudotypes encoding luciferase and harboring the indicated spike variants were incubated with serial dilutions of plasma for 1 hour at 37°C and then used to infect 293T-ACE2. Infection was measured by quantitating luciferase activity 72 hours post-infection. Neutralization half maximal inhibitory serum dilution (ID50) for the sera from vaccinated or convalescent individuals were determined using a normalized non-linear regression using GraphPad Prism (Table S6). Limit of detection is indicated by a dotted line (ID50=50). Distributions across replicates are represented by boxplots with a central median value and whiskers showing the 1.5x inter-quartile range. Significant group differences (from Welch’s one-way ANOVA with Tukey’s post hoc testing) are indicated using brackets and asterisks (* <0.05, ** < 0.01, *** < 0.001).

## Discussion

We have identified a highly divergent lineage of SARS-CoV-2 with evidence of host adaptation and unsustained deer-to-human transmission. WTD present many attributes important for reservoir sustainability including social behaviour, high density, highly transient populations with significant human-deer interfaces and sylvatic interfaces with other wildlife. A stable reservoir in WTD creates the potential for spillover into human and sylvatic wildlife populations over a broad geographic distance. In contrast to domestic mink, onward transmission mitigation among WTD is more challenging than for farmed species.

Phylogenetic analysis revealed that the Ontario WTD lineage shares a relatively recent common ancestor with mink- and human-derived viral sequences from nearby Michigan. This includes specific mutational similarities such as a subset of mink sequences from Michigan exhibiting a rare 12-nucleotide insertion in the ORF1a gene that was also present in the Ontario WTD lineage. Two S gene mutations, F486L and N501T have been associated with mustelid (mink or ferret) host adaptations, and N501T has been associated with enhanced ACE2 binding and entry into human (Huh7) cells ^7,18,37,38^. Notably, the Ontario WTD SARS-CoV-2 genomes did not harbour the relatively well-described S:Y453F mutation associated with mink and increased replication and morbidity in ferrets, but reduced replication in primary human airway epithelial cells ^39^. Many of the mutations in the divergent Ontario WTD SARS-CoV-2 genomes have not been described previously or are infrequent and uncharacterized. These WTD genomes provide new insights into viral evolution and inferred virus mobility in animal species outside of the human population.

The mutational spectra of SARS-CoV-2 genomes from WTD, mink, and humans vary between hosts, as highlighted by significant differences within Clade 20C. Importantly, this provides evidence supporting the hypothesis that mutational spectra can be used to infer viral host-species ^9,40^. Furthermore, the frequency of C>U and A>G mutations differed between hosts, which may reflect host cell activity, such as restriction factors (e.g. apolipoprotein B mRNA editing enzyme, catalytic polypeptide-like or APOBEC family of mRNA editing proteins), RNA editing enzymes (ADAR1), and/or reactive oxygen species ^41–44^. We also observed that the mutational spectrum of the Ontario WTD (and human) sequences is similar to that of other deer. However, this observation does not imply that mutations found in these sequences are related to those found in other deer. Rather, it is likely that the interaction between the viral genome and various host factors will alter the mutation spectra in broad, host-specific ways ^9,40^. When placed in the context of the broader literature, our results provide further evidence this lineage of SARS-CoV-2 likely evolved in deer over time.

The absence of detectable positive selection in the WTD lineage, evidence of relaxation of selection within ORF1ab, and the distribution of mutations across the genome contrasts with the signatures of strong selection in the equivalently divergent Omicron VOC. While additional complete WTD genomes could enable a more nuanced analysis (e.g., whole-genome compartmental models and exclusion of terminal branches to limit within-host biases ^45^) it is clear that the evolutionary forces acting within the WTD lineage are considerably different to those in Omicron. From these results, and a phylogenetically distant most recent common ancestor from 2020, we can infer that the WTD lineage likely diverged in 2020 and has been maintained in wildlife under minimal selection pressure since then. It is possible that the absence of pre-existing host immunity permitted genetic drift to drive accumulation of neutral mutations (in combination with accumulation of mutations associated with animal adaptation).

It is unclear if the initial spillover occurred directly from humans to deer, or if an intermediate host such as mink or other yet undefined species was involved. The long branch length and period of unsampled evolution provide a number of possible scenarios (Fig. 6). The human sample in the Ontario WTD clade lies within a relatively small number of deer samples, making determining the exact relationship between the human- and WTD-derived viruses challenging (78% UFB). This clade could represent a spillover into WTD with a human spillback or the emergence of a virus reservoir in a wildlife species infecting both human and WTD. However, the epidemiological data, evidence of infectious virus from deer, and the paucity of SARS-CoV-2 surveillance in WTD relative to human cases suggest spillover in WTD followed by deer-to-human transmission is the most likely scenario (Fig. 6: Scenario 1).

**Fig. 6:**
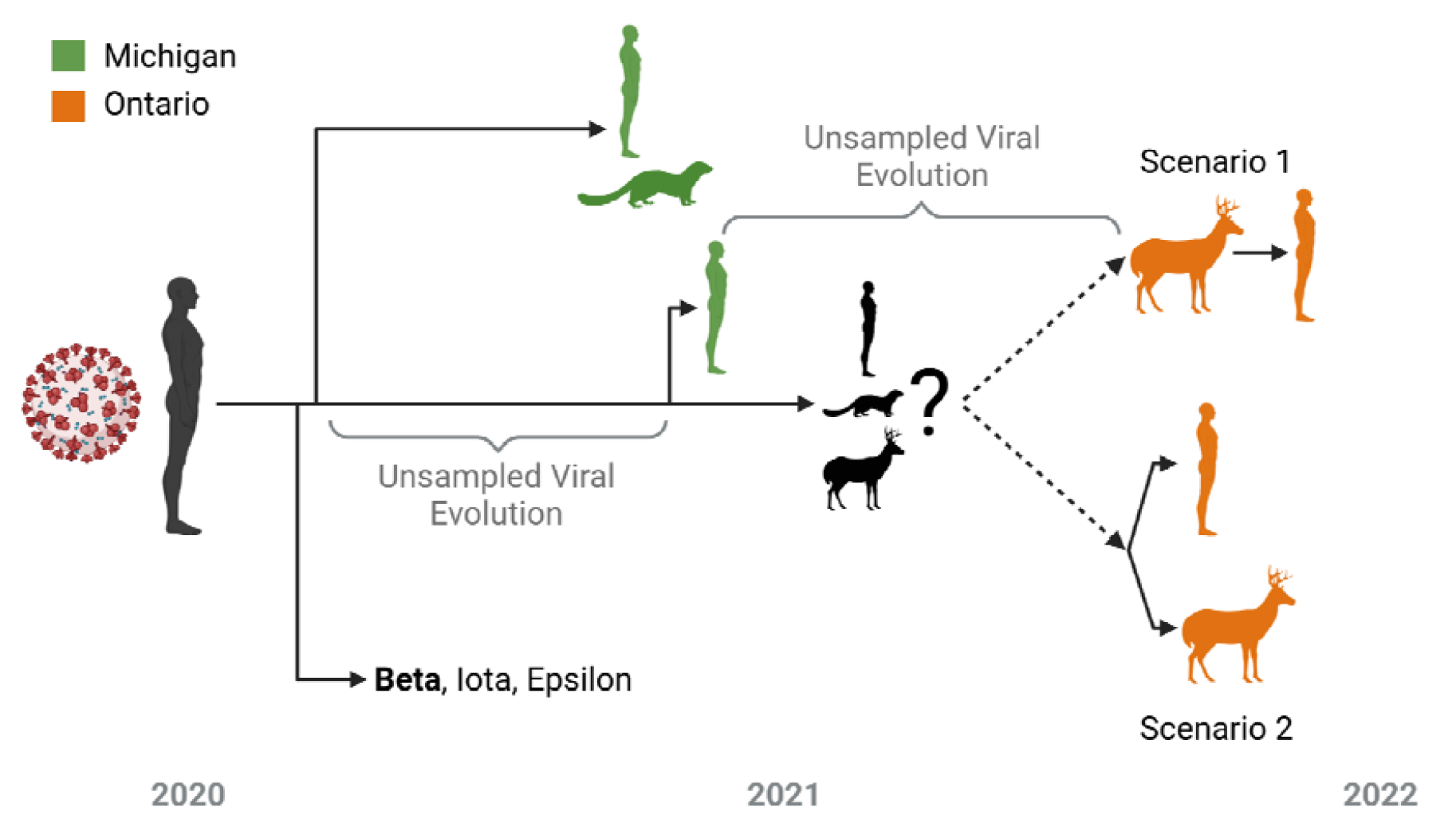
Overview of potential zoonotic scenarios underpinning the evolution of the Ontario WTD clade (and associated human case). The timeline and approximate relationship between the Beta VOC (bold), Iota/Epsilon VUIs, and viral samples in WTD, humans, and mink from both Michigan (green) and Ontario (orange) are displayed. As it likely emerged during one of the indicated poorly sampled periods of viral evolution, it is unclear whether the viral ancestor of the Ontario WTD clade was from an unknown animal (e.g., mink, WTD, or other species) or human reservoir. From this ancestor, there was either a spillback transmission from WTD to human (Scenario 1) or the emergence of a virus infecting both human and WTD (Scenario 2).

At this time, there is no evidence of recurrent deer-to-human or sustained human-to-human transmission of the Ontario WTD SARS-CoV-2 clade. However, there has been considerable reduction in human and WTD testing and genomic surveillance since the study samples were collected and we cannot exclude the possibility of sustained transmission within or between these host populations. Enhanced surveillance is critical given human population density and mobility in the region, coupled with WTD population dynamics. In addition, rapid characterization of this new lineage from biological and epidemiological perspectives is critical to understanding viral transmission, immune evasion, and disease in both wildlife and humans. Therefore, we assessed the ability of antibodies elicited following vaccination or infection to recognize and neutralize S and found that the mutations in the Ontario WTD lineage do not have significant impact on S antigenicity (acknowledging the limited number of plasma tested) (Fig. 5). More work is needed to determine the potential roles of the mutations on spike functions, and to understand the pathogenesis and transmission phenotypes of this virus.

Secondary wildlife reservoirs have the potential to fundamentally alter the ecology of SARS-CoV-2. Our work underscores the need for a broad international One Health focus to identify new intermediate or reservoir hosts capable of driving sustained transmission and divergent viral evolution. An examination of human drivers of spillover and spillback and knock-on effects on wildlife and human health is urgently needed to identify, develop and implement mitigation strategies, beginning with reducing viral activity in humans.

## Methods

### Deer sample collection and study area

Between November 1 and December 31, 2021, adult and yearling free-ranging WTD were sampled as part of the Ontario Ministry of Northern Development, Mines, Natural Resources and Forestry’s (NDMNRF) annual Chronic Wasting Disease (CWD) surveillance program. Samples were collected from hunter-harvested deer in Southwestern and Eastern Ontario and included nasal swabs and retropharyngeal lymph nodes (RPLNs). All samples were collected by staff who wore a mask and disposable gloves while sampling. Nasal swabs were stored in individual 2mL tubes with ∼1mL of universal transport media (UTM; Sunnybrook Research Institute) and RPLN tissues were stored dry in 2 mL tubes. After collection, samples were immediately chilled on ice packs then transferred to a -20°C freezer where they were held for up to one week. Samples were then transferred to a -80°C freezer where they were held until analysis. Location and date of harvest and demographic data (age/sex) were recorded for each animal when available.

### PCR screening and detection

RNA extractions and PCR testing of samples collected from deer were performed at the Sunnybrook Research Institute (SRI) in Toronto, Ontario. RNA extractions were conducted with 140 µL of nasal swab sample spiked with Armored RNA enterovirus (Asuragen; https://www.asuragen.com) via the Nuclisens EasyMag using Generic Protocol 2.0.1 (bioMérieux Canada Inc., St-Laurent, QC, Canada) according to manufacturer’s instructions; RNA was eluted in 50 µL. Tissue samples were thawed, weighed, minced with a scalpel, and homogenized in 600 µL of lysis buffer using the Next Advance Bullet Blender (Next Advance, Troy, NY USA) and a 5mm stainless steel bead at 5 m/s for 3 minutes. RNA from 30 mg tissue samples were extracted using Specific Protocol B 2.0.1 via Nuclisens EasyMag; RNA was eluted in 50 µL. Reverse-transcription polymerase chain reaction (RT-PCR) was performed using the Luna Universal Probe One-Step RT-qPCR kit (New England BioLabs; https://www.neb.ca). The 5 ‘ untranslated region (UTR) and the envelope (E) gene were used for SARS-CoV-2 RNA detection ^46^. Quantstudio 3 software (Thermo Fisher Scientific; https://www.thermofisher.com) was used to determine the cycle threshold (Ct). All samples were run in duplicate and samples with Ct<40 for both gene targets and Armored RNA enterovirus in at least one replicate were considered presumed positive. For tissue samples, the presence of inhibitors was assessed by a 1:5 dilution of one of the replicates. Samples were considered inconclusive if no Armored enterovirus was detected or if only one gene target was detected and were re-extracted for additional analysis. Samples were considered indeterminate if inconclusive after re-extraction or if no original material was left. Presumed positive samples were further analyzed for human RNAse P to rule out potential human contamination ^11^.

Original material from presumed positive samples detected at SRI were sent to the Canadian Food Inspection Agency (CFIA) for confirmatory PCR testing. The MagMax CORE Nucleic Acid Purification Kit (ThermoFisher Scientific) and the automated KingFisher Duo Prime magnetic extraction system was used to extract total RNA spiked with Armored RNA enterovirus. The enteroviral armored RNA was used as an exogenous extraction control. The E and nucleocapsid (N) genes were used for confirmatory SARS-CoV-2 RNA detection ^7^. Master mix for qRT-PCR was prepared using TaqMan Fast Virus 1-step Master Mix (ThermoFisher Scientific) according to the manufacturer’s instructions. Reaction conditions were 50°C for 5 minutes, 95°C for 20 seconds, and 40 cycles of 95°C for 3 seconds then 60°C for 30 seconds. Runs were performed by using a 7500 Fast Real-Time PCR System (Thermofisher, ABI). Samples with Ct <36 for both gene targets were considered positive.

### WGS sequencing

WGS was performed at both SRI and CFIA using independent extractions and sequencing methods. At SRI, DNA was synthesized from extracted RNA using 4 μL LunaScript RT SuperMix 5X (New England Biolabs, NEB, USA) and 8 μL nuclease free water, were added to 8 μL extracted RNA. cDNA synthesis was performed under the following conditions: 25 °C for 2 min, 55 °C for 20 min, 95 °C for 1 min, and holding at 4 °C.

The ARTIC V4 primer pool (https://github.com/artic-network/artic-ncov2019) was used to generate amplicons from the cDNA. Specifically, two multiplex PCR tiling reactions were prepared by combining 2.5 μL cDNA with 12.5 μL Q5 High-Fidelity 2X Master Mix (NEB, USA), 6μL nuclease free water, and 4 μL of respective 10 μM ARTIC V4 primer pool (Integrated DNA Technologies). PCR cycling was then performed in the following manner: 98 °C for 30 s followed by 35 cycles of 98 °C for 15 s and 63 °C for 5 min.

PCR reactions were then both combined and cleaned using 1X ratio Sample Purification Beads (Illumina) at a 1:1 bead to sample ratio. The amplicons were quantified using the Qubit 4.0 fluorometer using the 1X dsDNA HS Assay Kit (Thermo Fisher Scientific, USA) and sequencing libraries prepared using the Nextera DNA Flex Prep kit (Illumina, USA) as per manufacturer’s instructions. Paired-end (2×150 bp) sequencing was performed on a MiniSeq with a 300–cycle reagent kit (Illumina, USA) with a negative control library with no input SARS-CoV-2 RNA extract.

WGS performed at CFIA used extracted nucleic acid quantified using the Qubit™ RNA High Sensitivity (HS) Assay Kit on a Qubit™ Flex Fluorometer (Thermo Fisher Scientific). 11uL or 200ng of total RNA was subject to DNase treatment using the ezDNase™ Enzyme (Thermo Fisher Scientific) according to the manufacturer’s instructions. DNase treated RNA was then used for library preparation and target sequence capture according to the ONETest™ Coronaviruses Plus Assay protocol (Fusion Genomics ^47^). The enriched libraries were then quantified using the Qubit™ 1x dsDNA HS Assay Kit on a Qubit™ Flex Fluorometer (Thermo Fisher Scientific) and subsequently pooled in equimolar amounts prior to fragment analysis on 4200 TapeStation System using the D5000 ScreenTape Assay (Agilent). The final pooled library was sequenced an Illumina MiSeq using a V3 flowcell and 600 cycle kit (Illumina).

Human specimens are received at Public Health Ontario Laboratory (PHOL) for routine SARS-CoV-2 diagnostic testing (RT-PCR) from multiple healthcare settings across the province, including hospitals, clinics and COVID-19 assessment centres. The human sample (ON-PHL-21-44225) was sequenced at PHOL using an Illumina-based ARTIC V4 protocol (10.17504/protocols.io.b5ftq3nn), similar to the deer sequencing methods. Briefly, cDNA was synthesized using LunaScript reverse transcriptase (New England BioLabs). Amplicons were generated with premixed ARTIC V4 primer pools (Integrated DNA Technologies). Amplicons from the two pools were combined, purified with AMPure XP beads (Beckman Coulter) and quantified. Genomic libraries were prepared using the Nextera XT DNA Library Preparation Kit (Illumina, San Diego, CA) and genomes were sequenced as paired-end (2 × 150-bp) reads on an Illumina MiSeq instrument.

### Genomic analysis

Paired-end illumina reads from ARTIC V4 and Fusion Genomics sequencing were initially analysed separately with the nf-core/viralrecon Nextflow workflow (v2.3) ^48–50^ which ran: FASTQC (v0.11.9) ^51^ read-level quality control, fastp (v0.20.1) ^52^ quality filtering and adapter trimming, Bowtie2 (v2.4.2) ^53^ read mapping to Wuhan-Hu-1 (MN908947.3) ^54^ SARS-CoV-2 reference, Mosdepth (v0.3.1)^55^/Samtools (v.1.12) ^56^ read mapping statistics calculation, iVar (v1.3.1) ^57^ ARTIC V4 primer trimming, variant calling, and consensus generation; SnpEff (v5.0) ^58^/SnpSift (v4.3t) ^59^ for variant effect prediction and annotation; and Pangolin (v3.1.20) ^60^ with PangoLEARN (2022-01-05), Scorpio (v0.3.16) ^61^, and Constellations (v.0.1.1) was used for PANGO lineage ^62^ assignment. iVar primer trimmed soft-clipped read alignments were converted to hard-clipped alignments with fgbio ClipBam (http://fulcrumgenomics.github.io/fgbio/). Reads from hard-clipped BAM files were output to FASTQ files with “samtools fastq”. nf-core/viralrecon was re-run in amplicon mode without iVar primer trimming on the combined Fusion Genomics and ARTIC V4 primer trimmed FASTQ reads to generate the variant calling, variant effect and consensus sequence results used in downstream analyses. Additional quality control steps to check for negative control validity, drop-out, sample cross-contamination, and excess ambiguity were performed using ncov-tools v1.8.0 ^63^. The mutations identified by the Nextclade (v1.10.2) ^64^ with 2022-01-05 database and xlavir (v0.6.1) report were manually searched in outbreak.info’s “Lineage | Mutation Tracker” (on 2022-02-02) ^65^ to get information on the prevalence of observed mutations globally and within Canada. Mutations were also investigated for presence in specific lineages including VOCs, Michigan mink samples, and other animal samples. Finally, mutations were searched in GISAID (on 2022-02-02) to tally the number of non-human hosts each mutation had been observed in.

It should be noted there were some limitations in genome quality and coverage that may have resulted in failure to detect additional mutations that were present. All Ontario WTD clade samples (including the associated human case) had missing terminal domains and contained internal regions with no or low coverage when sequenced using the ARTIC v4 amplicon scheme. This is a widespread issue that may explain the rarity of the 3 ‘ proximal ORF10:L37F in GISAID. Significantly in our samples this meant there was no or <10x coverage in all 5 WTD sequences from ∼27000-27177 (dropout of ARTICv4 amplicons 90-91) which includes regions of the M gene. However, by combining the ARTIC v4 sequencing with additional sequencing using probe-based enrichment we were able to compensate for this dropout and generate high coverage and completeness (<100 positions with no coverage in all WTD and <100 positions with <10X coverage in 3/5 WTD genomes; see Table S7).

### Phylogenetics

To evaluate possible sampling biases due to the poorly defined and diverse B.1 and B.1.311 lineages and select closely related publicly available sequences for further phylogenetic analysis, an UShER (https://genome.ucsc.edu/cgi-bin/hgPhyloPlace) ^33^ based phylogenetic placement analysis was performed using the 7,217,299 sample tree (derived from UShER placement of GISAID, GenBank, COG-UK, and CNCB onto 13-11-20 sarscov2phylo ML tree) via the SHUShER web-portal (shusher.gi.ucsc.edu). Phylogenetic analyses were performed using CFIA-NCFAD/scovtree Nextflow workflow (v1.6.0) (https://github.com/CFIA-NCFAD/scovtree/) with the consensus sequences contextualised with closely related sequences identified by UShER and randomly sampled representative sequences from major WHO SARS-CoV-2 clades from GISAID ^13^ (downloaded 2022-02-10). This workflow generated a multiple sequence alignment using Nextalign CLI (v1.10.1) ^64^ and inferred a maximum-likelihood (ML) phylogeny using IQ-TREE (v2.2.0_beta) ^66^ using the GTR model for visualisation with Phylocanvas ^67^ via shiptv (v0.4.1) (https://github.com/CFIA-NCFAD/shiptv) and ggtree ^68^. Divergence times (and ancestral mutation states) were inferred from this phylogeny via treetime (v0.8.6) (^69^ and augur (v14.0.0) ^70^ using a scalar optimized coalescent model and a clock rate of 0.0009+/-0.0006 and 8 interquartile-distance filter. These clock values were derived from ordinary least squares regression of root to tip divergence distances using statsmodel (v0.13.2) ^71^ and ETE3 ^72^ (v3.1.2) phylogenetic library.

A subset of 157 taxa from an ancestral clade of the Ontario WTD and human clade were selected from the global phylogenetic tree shown in Fig. 2 to generate the phylogenetic tree shown in Fig. 3. Multiple sequence alignment of this subset of sequences was performed with MAFFT (v7.490) ^73^. A maximum-likelihood phylogenetic tree was inferred with IQ-TREE (v2.2.0_beta) using the GTR model and 1000 ultrafast bootstrap replicates ^74^. Nextclade (v1.10.2) analysis was used to determine amino acid mutations and missing or low/no coverage regions from the sample genome sequences. Amino acid mutation profiles were determined relative to the Ontario WTD and human samples, discarding mutations that were not present in any of the Ontario samples. Taxa with duplicated amino acid mutation profiles were pruned from the tree, keeping only the first observed taxa with a duplicated profile.

Recombination analyses were performed using 3Seq (v1.7) ^29^ and Bolotie (e039c01) ^30^. Specifically, 3Seq was executed with WTD+Human sequences and the most recent example of each lineage found in Canada and closest samples in GISAID in subtree (n=595). Bolotie was executed using the WTD+Human sequences and two datasets, the provided pre-computed 87,695 probability matrix and a subsample of the earliest and latest example of each lineage in GISAID with all animal-derived samples and closest usher samples (n=4,688). Additionally, HyPhy’s ^31^ (v2.5.31) Genetic Algorithm Recombination Detection (GARD) method ^32^ was applied to local alignment and ML phylogeny (Fig. 3) for all possible sites. Phylogenies were inferred using IQTree for segments either side of the identified putative breakpoint and the ON WTD clade and local topology was unchanged. Sequence statistics such as C>T rate were directly calculated from nextclade results (v1.10.2 with 2022-01-05 database). Additional figures were generated and annotated using BioRender ^75^ and Inkscape ^76^.

A phylogenetic approach with HyPhy ^31^ (v2.5.31) was used to investigate signatures of selection within the Ontario WTD clade relative to the wider B.1 background lineage. To ensure necessary genomic completeness for codon alignment, all B.1 sequences in GISAID (as of 2022-03-08) were filtered to those <0.1% N’s with full date information. Genomes with 0% N were removed to avoid biases from consensus workflows which replace undetermined sequence with reference genome. From this, all animal-derived (49 mink, 1 cat) sequences and 100 randomly sampled human B.1 sequences were extracted (n=150). Finally, the WH0-1 reference genome was added to this alignment along with the 5 complete ON WTD genomes, associated human sequence, and the 2 most closely related Michigan human samples (MI-MDHSS-SC23517, M-MDHSS-SC22669). Virulign ^77^ (v1.0.1) was then used to generate codon alignments for E, M, N, S, ORF1ab, ORF3a, ORF6, ORF7a, ORF7b, ORF8, and ORF10 genes relative to the Wuhan-Hu-1 (MN908947.3) reference. ML-phylogenies were inferred for these alignments using raxml-ng (v1.0.2) ^78^ with the GTR model and 3 parsimony-based starting trees. These phylogenies were manually inspected, rooted on Hu-1, and the Ontario WTD clade branches labeled using phylowidget ^36,79^. Genes for which the phylogeny did not have a resolved WTD clade (ORF7a and ORF7b) or a viable codon alignment without any internal stop codons (ORF8 and ORF10) were excluded from further analyses. For each gene, signatures of positive selection were evaluated using HyPhy’s adaptive branch-site random effects likelihood (aBSREL) method ^35^ and signatures of gene-wide episodic diversification were evaluated using the Branch-Site Unrestricted Statistical Test for Episodic Diversification (BUSTED) method ^36^ with 10 starting points. Finally, evidence of intensification or relaxation of selection was investigated using the RELAX method ^80^ with 10 starting points and synonymous rate variation. Additional code for divergence dating, recombination, and selection analyses can be found under DOI: 10.5281/zenodo.6533999.

### Analysis of mutational spectrum

The mutational spectra was created using a subset of 3,645 sequences used to create the high-quality global phylogeny. Included in this dataset are a collection of seven unique WTD samples from Ontario (samples 4538, 4534, 4662, 4649, 4581, 4645, and 4658; the 5 high quality genomes plus 2 genomes with lower coverage), three WTD samples from Quebec (samples 4055, 4022, 4249), and one human sample from Ontario (ON-PHL-21-44225). The counts for each type of nucleotide change, with respect to the reference strain, were used to create a 12-dimensional vector. The final dataset consisted of all human, mink, and deer 20 samples originating from the United States of America and Canada with at least 15 mutations. The counts were converted into the mutation spectrum by simply dividing each count by the sum of the counts in each sample ^13^. To investigate differences in mutation spectra between hosts, a distance-based Welch MANOVA (dbWMANOVA) was run ^81^. If a significant difference was detected, a pairwise distance-based Welch t-Test was used to identify which pair of hosts significantly differed ^82^. These tests were used since they are more robust on unbalanced and heteroscedastic data ^81,82^.

### Codon usage analysis

Consensus sequences of SARS-CoV-2 samples from this and previous studies and additional sequences gathered from public databases were used. The sequences include the reference SARS-CoV-2 Wuhan-Hu-1 (NCBI NC_045512), SARS-CoV-2 mink/Canada/D/2020 (GISAID EPI_ISL_717717), SARS-CoV-2 mink/USA/MI-20-028629-004/2020 (GISAID EPI_ISL_2834697), *Cervid atadenovirus A* 20-5608 (NCBI OM470968) ^83^, EHDV serotype 2 / strain Alberta (NCBI AM744997 - AM745006), Epizootic Hemorrhagic Disease Virus, EHDV serotype 1 / New Jersey (NCBI NC_013396 - NC_013405), EHDV 6 isolate OV208 (NCBI MG886400 - MG886409), and Elk circovirus Banff/2019 (NCBI MN585201) ^84^ were imported into Geneious (v.9.1.8) ^85^. Annotations for the coding sequences of SARS-CoV-2 samples were transferred from the reference sequence SARS-CoV-2 Wuhan-Hu-1 (NC_ 045512) using the Annotate from Database tool. The coding sequences were extracted using the Extract Annotations tool for all viral sequences. An annotated file of the coding sequences for the *Odocoileus virginianus texanus* isolate animal Pink-7 (GCF_002102435.1) genome was downloaded from NCBI (https://ftp.ncbi.nlm.nih.gov/genomes/all/annotation_releases/9880/100/GCF_002102435.1_Ovir.te_1.0/). Coding sequences were input into CodonW (http://codonw.sourceforge.net/) with settings set to concatenate genes and output to file set to codon usage. Codon usage indices were set to the effective number of codons (ENc), GC content of gene (G+C), GC of silent 3rd codon position (GC3s), silent base composition, number of synonymous codons (L_sym), total number of amino acids (L_aa), hydrophobicity of protein (Hydro), and aromaticity of protein (Aromo).

### Virus isolation

For virus isolation, T25 flasks were seeded to confluency one day prior to infection with cathepsin L knock out VeroE6 cells overexpressing TMPRSS2. The following day swab samples were vortexed and spun down and 200 ul of the swab samples medium was combined with 16 ug/ml working concentration of TPCK-treated trypsin (New England Biolabs Ltd), 2x A/A/p/s antifungal/antibiotic solution (Wisent), and a 0.1% working concentration of BSA (Thermo Fisher Scientific) and added to the cell monolayer after removal of the medium. Samples were subjected to a 45 minute adsorption with rocking every 5 minutes after which the inoculum was removed and discarded and the monolayer was either washed once with 2 ml of D1 to remove blood cells present in the samples (4581, 4649, and 4676) or not washed (4645, 4658, 4662) and 5 ml of DMEM medium with 1% FBS and antibiotics was added to the flask and incubated at 37°C with 5% CO_2_. At 4 dpi, samples with visible cytopathic effect (CPE) (partial, 50% or less rounded or detached cells) were harvested followed by collection and centrifugation at 4000 xg for 10 minutes at 20°C. The harvested supernatants were aliquoted and stored at -80°C or inactivated and removed from the CL3 laboratory and RNA was extracted with the QIAamp^®^ Viral RNA Mini kit (Qiagen), and stored at -20°C until downstream analyses were carried out. All infectious work was performed under biosafety level 3 (BSL-3) conditions.

### Codon-optimized Spike constructs, cells, sera and antibodies

Expression constructs of S mutants corresponding to samples 4581/4645 (S:H49Y, S:T95I, S:Δ143-145InsD, S:F486L, S:N501T, S:D614G), 4658 (S:T22I, S:H49Y, S:T95I, S:Δ143-145InsD, S:S247G, S:F486L, S:N501T, S:D614G) and ON-PHL-21-44225 (S:H49Y, S:T95I, S:Δ143-145InsD, S:F486L, S:N501T, S:Q613H, S:D614G) were generated by overlapping PCR as described previously ^86^. S:D614G and S Omicron (BA.1) constructs were described elsewhere^87^. All constructs were cloned in pCAGGS and verified by Sanger sequencing.

HEK293T cells (ATCC) were cultured in Dulbecco’s Minimum Essential Medium (DMEM) supplemented with 10% fetal bovine serum (FBS, Sigma), 100 U/mL penicillin, 100 µg/mL streptomycin, and 0.3 mg/mL L-glutamine (Invitrogen) and maintained at 37°C, 5% CO_2_ and 100% relative humidity.

Serum samples were obtained from consenting participants in several cohort studies with sample collection and sharing for this analysis approved by the Sinai Health System Research Ethics Board (#22-0030-E). Plasma of SARS-CoV-2 naïve, naïve-vaccinated (28-40 days post two- or three-doses of BNT162b2), and unvaccinated SARS-CoV-2 Delta previously infected donors were collected (Table S8), heat-inactivated for 1 h at 56°C, aliquoted and stored at - 80°C until use. The conformation-independent monoclonal anti-S2 CV3-25 from a convalescent individual was described and produced as described previously ^88,89^. The goat anti-human IgG conjugated with Alexafluor-647 was purchased from Invitrogen (A21445).

### Spike binding assays

Hek293T cells seeded in a 10cm Petri dish at 70% confluency were transfected with 10µg of SARS-CoV-2 spike protein plasmid, 1µg of lentiviral vector bearing green fluorescent protein (GFP) (PLV-eGFP) (gift from Pantelis Tsoulfas, Addgene plasmid # 36083) ^90^ using Jetprime transfection reagent (Polyplus, cat#101000046) according to the manufacturer’s instructions. At 16 hours post transfection, the cells were stained with sera samples (1:250 dilution) for 45min at 37°C. Alexa Fluor-647-conjugated goat anti-human IgG (H+L) was used to detect plasma binding of the treated cells following 1 hour incubation at room temperature. Samples were washed once with PBS, fixed in 1% paraformaldehyde and acquired using BD LSR Fortessa Flow cytometer (BD Biosciences). The seropositivity threshold was defined based on: the mean of the MFIs for naïve samples plus three standard deviations. The data were normalized by surface expression based on the MFI of the monoclonal antibody CV3-25 (5ug/mL). The data analysis was performed using FlowJo 10.8.1. For each set of sera, binding was compared across samples using Welch’s (unequal variance) one-way ANOVA procedure and a post-hoc Tukey’s honestly significant difference test (using a family-wise error rate of 0.05) via the statsmodel library (v0.14.0) ^71^.

### Pseudotype production and neutralization assays

HEK293T seeded in 10-cm dishes were co-transfected with lentiviral packaging plasmid psPAX2 (gift from Didier Trono, Addgene #12260), lentiviral vector pLentipuro3/TO/V5-GW/EGFP-Firefly Luciferase (gift from Ethan Abela, addgene#119816), and plasmid encoding the indicated S construct at a 5:5:1 ratio using jetPRIME transfection reagent according to the manufacturer protocol. Twenty-four hours post-transfection, media were changed and supernatants containing lentiviral pseudotypes were harvested 48 h post-transfection, filtered with a 0.45 µM filter and stored at -80°C until use.

HEK293T stably expressing human ACE2 (293T-ACE2, kind gift of Hyeryun Choe, Scripps Research) were seeded in poly-D-lysine-coated 96-well plates. The next day, supernatants containing lentiviral pseudotypes were incubated with sera (serially diluted by five-fold, from 1:50 to 156,250) for 1 hour at 37° and then added to cells in the presence of 5 µg/mL polybrene. Seventy-two hours later, media were removed, and cells were rinsed in phosphate-buffered saline and lysed by the addition of 40 µl passive lysis buffer (Promega) followed by one freeze-thaw cycle. A Synergy Neo2 Multi-Mode plate reader (BioTek) was used to measure the luciferase activity of each well after the addition of 50-100 µl of reconstituted luciferase assay buffer (Promega) as per the manufacturer’s protocol. Neutralization half-maximal inhibitory dilutions (ID50) were calculated using Graphpad Prism and represent the plasma dilution that inhibits 50% of pseudotype transduction in 293T-ACE2. For each set of sera, neutralization was compared across samples using Welch’s (unequal variance) one-way ANOVA procedure and a post-hoc Tukey’s honestly significant difference test (using a family-wise error rate of 0.05) via the statsmodel library (v0.14.0) ^71^.

## Acknowledgments

We acknowledge the contributions of the Virus Detection, Molecular Diagnostics, DNA Core sections and the Biocomputing Centre of Public Health Ontario, and in particular Sarah Teatero for leading the genome sequencing efforts. We gratefully acknowledge contributions of SARS-CoV-2 genome sequences from other laboratories through GISAID (Table S9). We wish to thank the licensed Ontario deer hunters who submitted samples for wildlife disease surveillance, the staff of NDMNRF’s CWD surveillance program for their assistance with sample collection, and Sarah Hagey for GIS support. We also acknowledge the contributions of CFIA NCFAD’s Genomics Unit for their assistance with additional laboratory support and sequencing. S.M. and B.P. are members of the Canada and the Canadian Institutes for Health Research *Coronavirus Variants Rapid Response* Network (CoVaRR-Net). We are grateful to Dr. David Bulir from McMaster University for providing the primer and probe sequences for the 5 ‘ UTR gene SARS-CoV-2 PCR assay, and to Dr. Mikko Taipale for cathepsinL knockout TMPRSS2 overexpressing Vero cells. We appreciate the Public Health Agency of Canada for uploading the raw sequencing data for the Public Health Ontario human sequence to the Short Read Archive. The CV3-25 antibody was produced using the pTT vector kindly provided by the Canada Research Council. We also thank D. Leclair, J. Provencher, C. Soos, and A. Wilcox (Environment and Climate Change Canada) for comments on an earlier draft of the manuscript.

## Author Contribution Statement

Using the Contributor Roles Taxonomy - Conceptualisation: B.P., O.L., J.B., S.M., T.B; Methodology: F.M., O.L., J.K., B.P., D.S.; Software: F.M., P.K., O.L., J.S.; Formal analysis: F.M., P.K., J.G., O.L., J.S.; Investigation: M.C., K.N., P.A., G.L., A.A., B.V., J.B-S., H-Y.C, E.C., W.Y., M.G., M.S., M.P., G.S., A.M-A.; Data Curation: J.K., P.K., L.N., H.M.; Resources: A.M., E.A., A.F., G.G.; Writing - Original Draft: F.M., J.B., O.L., S.M., B.P., J.G.; Writing - Review & Editing: All authors; Visualisation: F.M., P.K., J.S.; Supervision: J.B., S.M., O.L.; Funding: B.P., S.M., M.C., J.K., O.L., J.B., T.B., L.N.

## Data and materials availability

All genomic sequence data are publicly available data through GISAID and SRA accession numbers are provided in the supplementary material (Table S1). All other data are available in the supplementary materials.

## Code availability

All computer code and analysis scripts used in the manuscript are archived at https://github.com/fmaguire/on_deer_spillback_analyses/ and can be referenced as DOI: 10.5281/zenodo.6533999.

## Extended Data

**Extended Data Figure 1:**
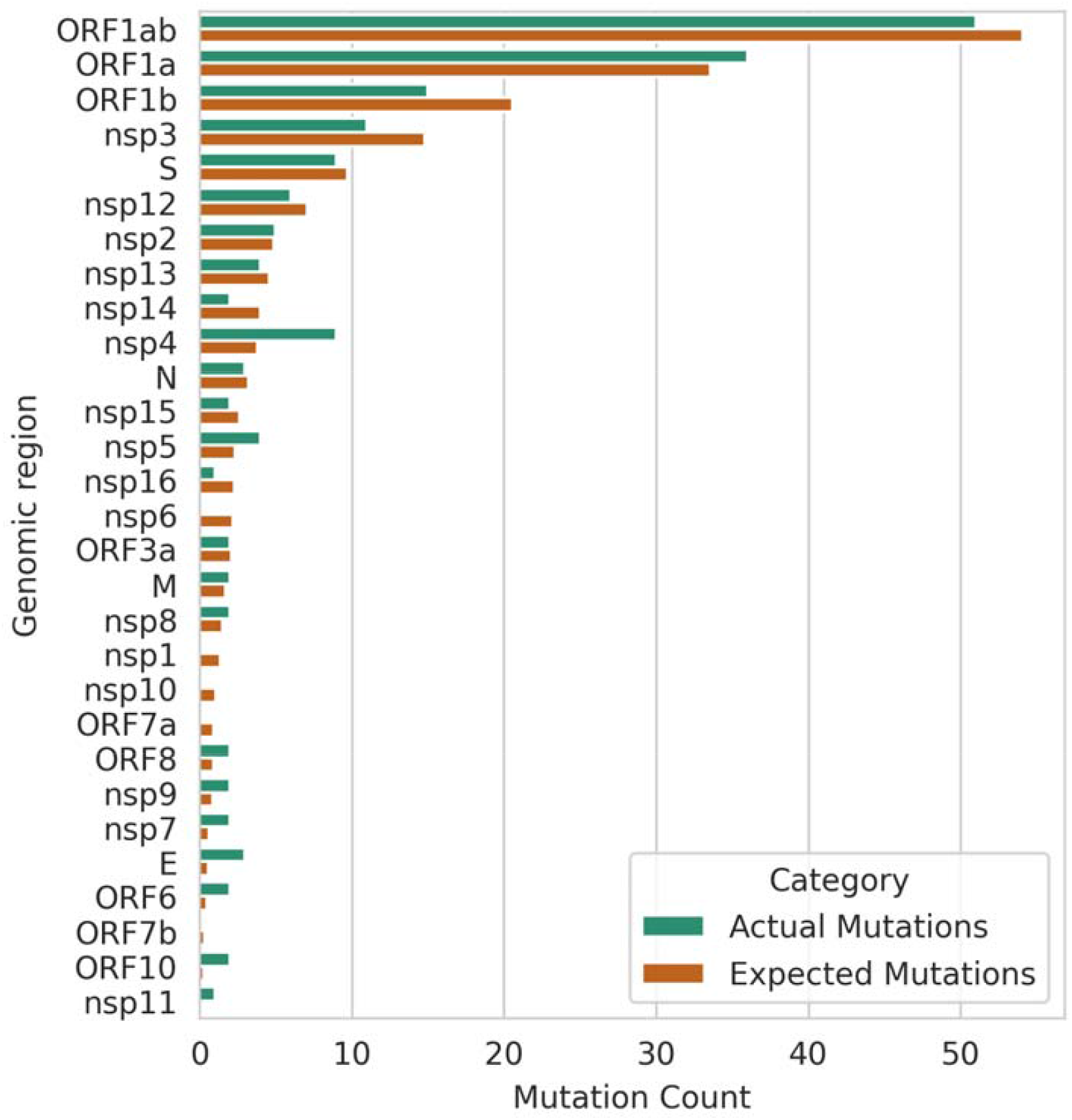
Observed vs expected counts within shared ON WTD clade mutations for each gene/product.

**Extended Data Figure 2:**
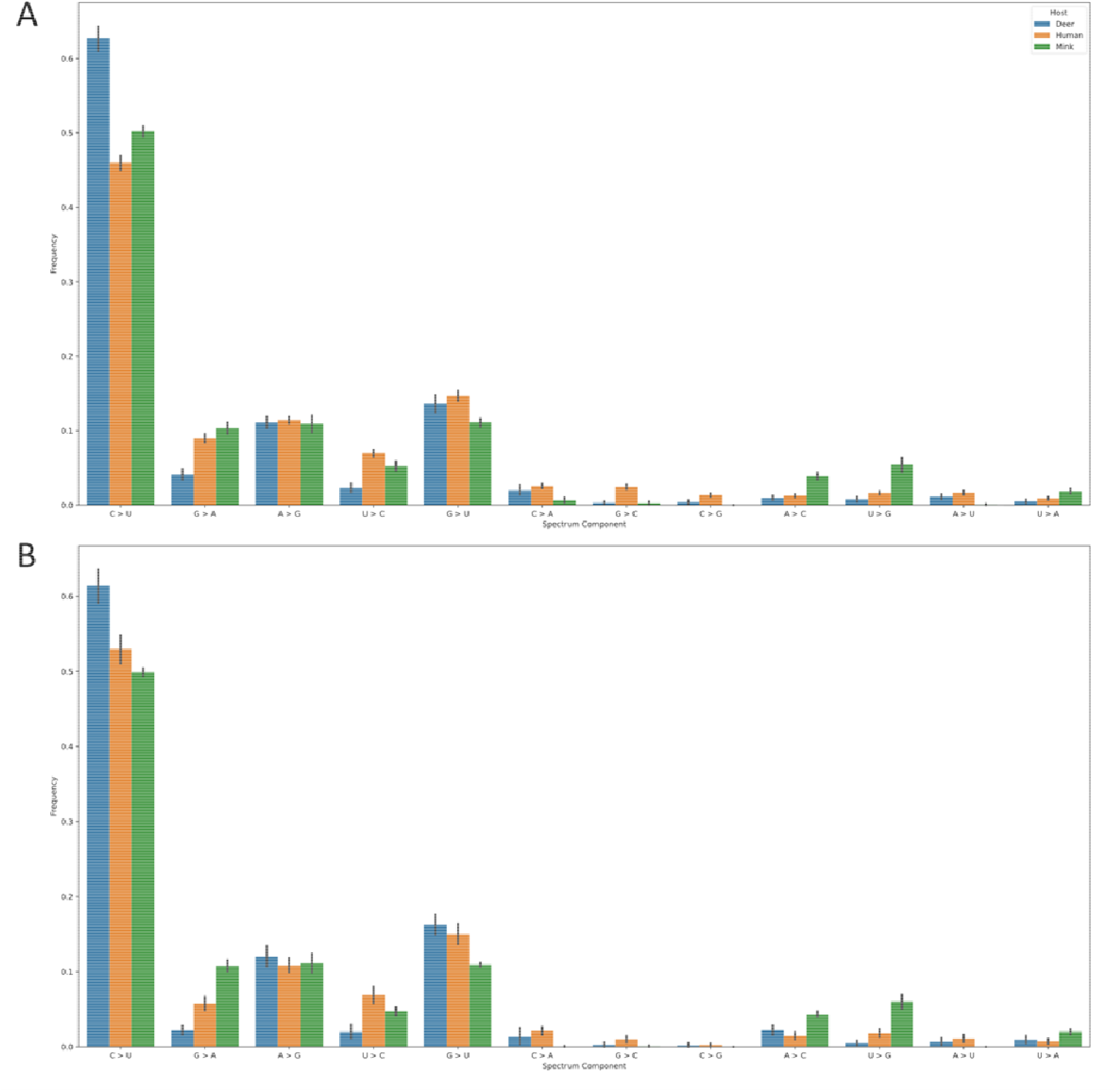
The frequency of different components of the mutational spectra from all samples (A) and only those found in Clade 20C (B).

**Extended Data Figure 3:**
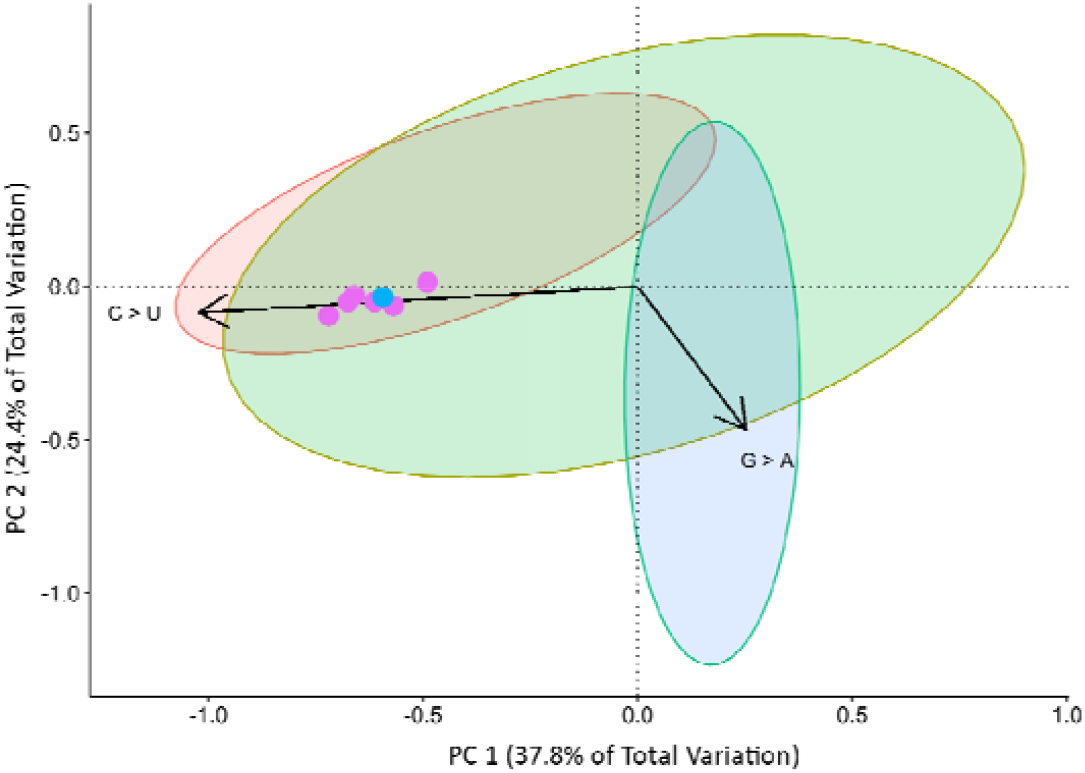
Principal Components Analysis of the mutational spectra of SARS-CoV-2 genomes isolated from different hosts within Clade 20C. The first two components account for 62.2% of the variation between the samples. Variation in the spectra along the first principal component associated with changes in the frequency of C>U. Samples appear to be spread along the first component by host-type. Ontario WTD samples (pink) and a human sample from Ontario (blue) appear close together in the projection, suggesting that they share a very similar mutation spectrum.

**Extended Data Figure 4:**
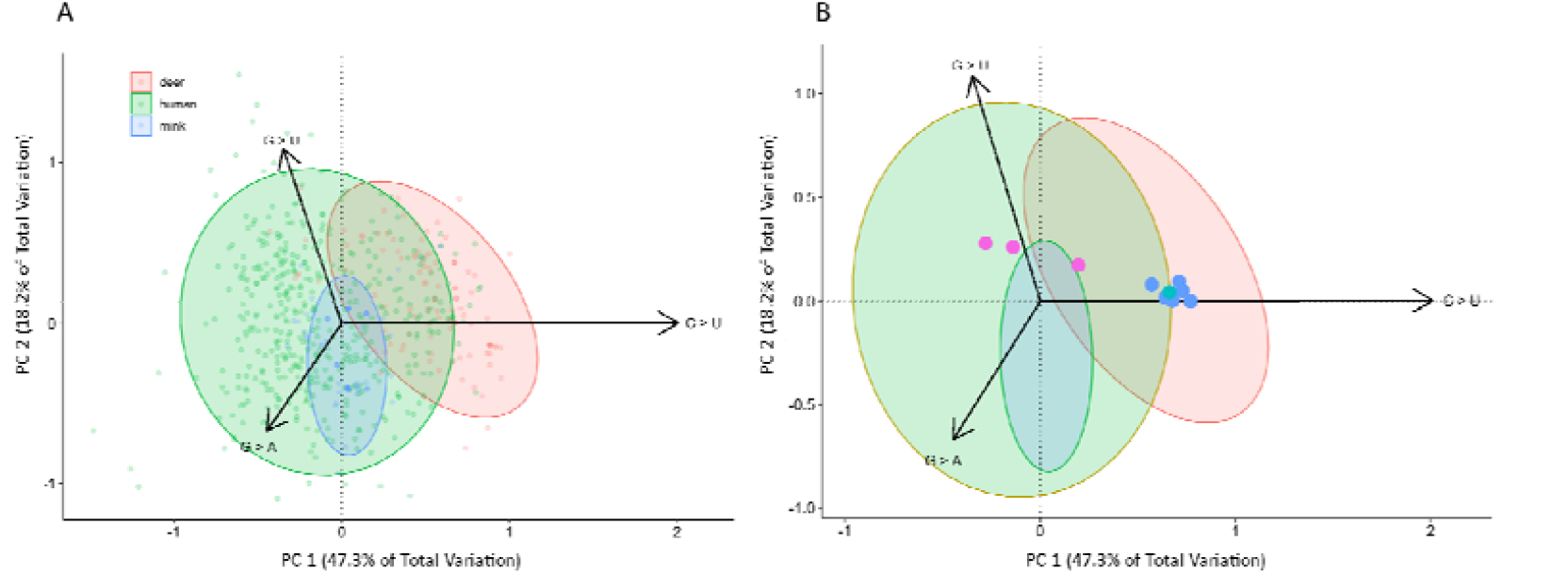
Principal Components Analysis of the mutational spectra from different SARS-CoV-2 variants. The first two components account for 65.5% of the variation between the samples. Variation in the spectra along the first principal component associated with changes in the frequency of C>U mutations. Interestingly, samples along this component are also differentiated by host-type. Variation along the second principal component reflect changes in the frequency of G>U and G>A mutations. (A) The PCA biplot showing a projection of all samples. (B) A simplified version of plot A highlighting the positions of WTD samples isolated from Ontario (Blue) and Quebec (Pink). A human isolate from Ontario appears in the same area of the plot (teal) as other Ontario WTD samples. Arrows are scaled to 30% of their original size to create a cleaner plot.

## Supplementary Material

**Figure S1: Usher based phylogenetic tree with Ontario WTD high quality (n=5) and partial (n=2) genomes and the 2022-02-21 USCS Usher build (shared in newick format)**.

Supplementary tables are shared in a Microsoft Excel workbook called “Supplementary_Tables.xlsx”. Table captions are provided below in the order in which they appear referenced in the manuscript and in the workbook. We have shared the Excel workbook as an auxiliary supplementary materials file.

**Table S1: Metadata associated with the 300 white-tailed deer and 1 human samples screened for SARS-CoV-2. Sample IDs used in the manuscript are the last 4 digits of the WILD-CoV Sample ID. * Samples not selected for sequencing with combined capture enrichment and ARTICv4 approach based on initial quality of ARTICv4 sequencing. + Human-derived sample sequenced by PHOL as part of public health surveillance. RPLN: Retrophyarngeal Lymph Nodes, RT-PCRs at SRI (UTR and E gene Ct<40) and CFIA (E and N2 gene Ct<36). Samples were considered positive (PVE), negative (NVE), or indeterminate (IND). For samples sent for confirmatory testing, positive (+) and negative (-) results are indicated in brackets from SRI and CFIA, respectively. Notably, results from SRI for two samples could only be considered inconclusive (+?) as remaining original material was depleted for confirmatory testing at CFIA**.

**Table S2: Summary of mutations within Ontario WTD clade (with associated human sequence) and their distribution across GISAID sequences, VOC, animal-derived viral sequences, and related Michigan mink sequences**.

**Table S3: Results of a distance-based Welch MANOVA investigating differences in the mutation spectrum between hosts within Nextstrain Clade 20C**.

**Table S4: Summary of codon usage bias analysis results across SARS-CoV-2 from WTD (including the Ontario lineage) and other cervid viruses**.

**Table S5: Plasma binding assay data (MFI normalised to CV3-25). Table S6: Neutralisation assay data (ID50)**.

**Table 7: Poor coverage regions in Ontario WTD clade (with associated human sequence) sequenced with combined ARTIC V4 and capture enrichment (ONETest) data**.

**Table S8: Description of the cohort and sera that was used for plasma binding and neutralisation assays**.

**Table S9: Acknowledgement table for sequences used from GISAID for phylogenetic and mutational signature analyses**.

## References

1. Andersen, K. G., Rambaut, A., Lipkin, W. I., Holmes, E. C. & Garry, R. F. The proximal origin of SARS-CoV-2. Nat. Med. 26, 450–452 (2020).

2. Boni, M. F. et al. Evolutionary origins of the SARS-CoV-2 sarbecovirus lineage responsible for the COVID-19 pandemic. Nat. Microbiol. 5, 1408–1417 (2020).

3. Cui, J., Li, F. & Shi, Z.-L. Origin and evolution of pathogenic coronaviruses. Nat. Rev. Microbiol. 17, 181–192 (2019).

4. Haagmans, B. L. et al. Middle East respiratory syndrome coronavirus in dromedary camels: an outbreak investigation. Lancet Infect. Dis. 14, 140–145 (2014).

5. Hu, B., Guo, H., Zhou, P. & Shi, Z.-L. Characteristics of SARS-CoV-2 and COVID-19. Nat. Rev. Microbiol. 19, 141–154 (2021).

6. Memish, Z. A. et al. Middle East respiratory syndrome coronavirus in bats, Saudi Arabia. Emerg. Infect. Dis. 19, 1819–1823 (2013).

7. Lu, L. et al. Adaptation, spread and transmission of SARS-CoV-2 in farmed minks and associated humans in the Netherlands. Nat. Commun. 12, 6802 (2021).

8. Mannar, D. et al. SARS-CoV-2 Omicron variant: Antibody evasion and cryo-EM structure of spike protein–ACE2 complex. Science (2022) doi:10.1126/science.abn7760.

9. Wei, C. et al. Evidence for a mouse origin of the SARS-CoV-2 Omicron variant. J. Genet. Genomics (2021) doi:10.1016/j.jgg.2021.12.003.

10. Yen, H.-L. et al. Transmission of SARS-CoV-2 delta variant (AY.127) from pet hamsters to humans, leading to onward human-to-human transmission: a case study. The Lancet 399, 1070–1078 (2022).

11. Hallmaier-Wacker, L. K., Munster, V. J. & Knauf, S. Disease reservoirs: from conceptual frameworks to applicable criteria. Emerg. Microbes Infect. 6, 1–5 (2017).

12. Abdel-Moneim, A. S. & Abdelwhab, E. M. Evidence for SARS-CoV-2 Infection of Animal Hosts. Pathogens 9, E529 (2020).

13. Shu, Y. & McCauley, J. GISAID: Global initiative on sharing all influenza data – from vision to reality. Eurosurveillance 22, 30494 (2017).

14. Tan, C. C. S. et al. Transmission of SARS-CoV-2 from humans to animals and potential host adaptation. bioRxiv 2020.11.16.384743 (2022) doi:10.1101/2020.11.16.384743.

15. Damas, J. et al. Broad host range of SARS-CoV-2 predicted by comparative and structural analysis of ACE2 in vertebrates. Proc. Natl. Acad. Sci. 117, 22311–22322 (2020).

16. Molenaar, R. J. et al. Clinical and Pathological Findings in SARS-CoV-2 Disease Outbreaks in Farmed Mink (Neovison vison). Vet. Pathol. 57, 653–657 (2020).

17. Shriner, S. A. et al. SARS-CoV-2 Exposure in Escaped Mink, Utah, USA. Emerg. Infect. Dis. J. 27, (2021).

18. Oude Munnink, B. B. et al. Transmission of SARS-CoV-2 on mink farms between humans and mink and back to humans. Science 371, 172–177 (2021).

19. Frutos, R. & Devaux, C. A. Mass culling of minks to protect the COVID-19 vaccines: is it rational? New Microbes New Infect. 38, 100816 (2020).

20. Pang, J. & Siu, T. Hong Kong to cull 2,000 hamsters after COVID-19 outbreak. Reuters (2022).

21. Peacock, T. P. et al. The SARS-CoV-2 variant, Omicron, shows rapid replication in human primary nasal epithelial cultures and efficiently uses the endosomal route of entry. 2021.12.31.474653 https://www.biorxiv.org/content/10.1101/2021.12.31.474653v1 (2022) doi:10.1101/2021.12.31.474653.

22. Shuai, H. et al. Emerging SARS-CoV-2 variants expand species tropism to murines. eBioMedicine 73, (2021).

23. Palmer, M. V. et al. Susceptibility of White-Tailed Deer (Odocoileus virginianus) to SARS-CoV-2. J. Virol. (2021) doi:10.1128/JVI.00083-21.

24. Chandler, J. C. et al. SARS-CoV-2 exposure in wild white-tailed deer (Odocoileus virginianus). Proc. Natl. Acad. Sci. 118, (2021).

25. Hale, V. L. et al. SARS-CoV-2 infection in free-ranging white-tailed deer. Nature 1–8 (2021) doi:10.1038/s41586-021-04353-x.

26. Kotwa, J. D. et al. First detection of SARS-CoV-2 infection in Canadian wildlife identified in free-ranging white-tailed deer (Odocoileus virginianus) from southern Québec, Canada. 2022.01.20.476458 https://www.biorxiv.org/content/10.1101/2022.01.20.476458v1 (2022) doi:10.1101/2022.01.20.476458.

27. Kuchipudi, S. V. et al. Multiple spillovers and onward transmission of SARS-CoV-2 in freeliving and captive white-tailed deer. 2021.10.31.466677 https://www.biorxiv.org/content/10.1101/2021.10.31.466677v2 (2021) doi:10.1101/2021.10.31.466677.

28. Marques, A. D. et al. Evolutionary Trajectories of SARS-CoV-2 Alpha and Delta Variants in White-Tailed Deer in Pennsylvania. 2022.02.17.22270679 https://www.medrxiv.org/content/10.1101/2022.02.17.22270679v2 (2022) doi:10.1101/2022.02.17.22270679.

29. Lam, H. M., Ratmann, O. & Boni, M. F. Improved Algorithmic Complexity for the 3SEQ Recombination Detection Algorithm. Mol. Biol. Evol. 35, 247–251 (2018).

30. Varabyou, A., Pockrandt, C., Salzberg, S. L. & Pertea, M. Rapid detection of inter-clade recombination in SARS-CoV-2 with Bolotie. Genetics 218, iyab074 (2021).

31. Kosakovsky Pond, S. L. et al. HyPhy 2.5—A Customizable Platform for Evolutionary Hypothesis Testing Using Phylogenies. Mol. Biol. Evol. 37, 295–299 (2020).

32. Kosakovsky Pond, S. L., Posada, D., Gravenor, M. B., Woelk, C. H. & Frost, S. D. W. Automated Phylogenetic Detection of Recombination Using a Genetic Algorithm. Mol. Biol. Evol. 23, 1891–1901 (2006).

33. Turakhia, Y. et al. Ultrafast Sample placement on Existing tRees (UShER) enables real-time phylogenetics for the SARS-CoV-2 pandemic. Nat. Genet. 53, 809–816 (2021).

34. Public Health Ontario. SARS-CoV-2 Whole Genome Sequencing in Ontario,. (2022).

35. Smith, M. D. et al. Less Is More: An Adaptive Branch-Site Random Effects Model for Efficient Detection of Episodic Diversifying Selection. Mol. Biol. Evol. 32, 1342–1353 (2015).

36. Murrell, B. et al. Gene-Wide Identification of Episodic Selection. Mol. Biol. Evol. 32, 1365–1371 (2015).

37. Starr, T. N. et al. Deep Mutational Scanning of SARS-CoV-2 Receptor Binding Domain Reveals Constraints on Folding and ACE2 Binding. Cell 182, 1295-1310.e20 (2020).

38. Han, P. et al. Molecular insights into receptor binding of recent emerging SARS-CoV-2 variants. Nat. Commun. 12, 6103 (2021).

39. Zhou, J. et al. Mutations that adapt SARS-CoV-2 to mink or ferret do not increase fitness in the human airway. Cell Rep. 38, (2022).

40. Shan, K.-J., Wei, C., Wang, Y., Huan, Q. & Qian, W. Host-specific asymmetric accumulation of mutation types reveals that the origin of SARS-CoV-2 is consistent with a natural process. Innov. N. Y. N 2, 100159 (2021).

41. De Maio, N. et al. Mutation Rates and Selection on Synonymous Mutations in SARS-CoV-2. Genome Biol. Evol. 13, evab087 (2021).

42. Mourier, T. et al. Host-directed editing of the SARS-CoV-2 genome. Biochem. Biophys. Res. Commun. 538, 35–39 (2021).

43. Ringlander, J. et al. Impact of ADAR-induced editing of minor viral RNA populations on replication and transmission of SARS-CoV-2. Proc. Natl. Acad. Sci. 119, (2022).

44. Simmonds, P. & Ansari, M. A. Extensive C->U transition biases in the genomes of a wide range of mammalian RNA viruses; potential associations with transcriptional mutations, damage-or host-mediated editing of viral RNA. PLOS Pathog. 17, e1009596 (2021).

45. Pond, S. L. K. et al. Adaptation to Different Human Populations by HIV-1 Revealed by Codon-Based Analyses. PLOS Comput. Biol. 2, e62 (2006).

46. LeBlanc, J. J. et al. Real-time PCR-based SARS-CoV-2 detection in Canadian laboratories. J. Clin. Virol. Off. Publ. Pan Am. Soc. Clin. Virol. 128, 104433 (2020).

47. Zhan, S. H. et al. Target capture sequencing of SARS-CoV-2 genomes using the ONETest Coronaviruses Plus. Diagn. Microbiol. Infect. Dis. 101, 115508 (2021).

48. Di Tommaso, P. et al. Nextflow enables reproducible computational workflows. Nat. Biotechnol. 35, 316–319 (2017).

49. Ewels, P. A. et al. The nf-core framework for community-curated bioinformatics pipelines. Nat. Biotechnol. 38, 276–278 (2020).

50. Patel, H. et al. nf-core/viralrecon: nf-core/viralrecon v2.3 - Copper Coatimundi. (Zenodo, 2022). doi:10.5281/zenodo.5974693.

51. Andrews, S. FastQC: a quality control tool for high throughput sequence data. (2010).

52. Chen, S., Zhou, Y., Chen, Y. & Gu, J. fastp: an ultra-fast all-in-one FASTQ preprocessor. Bioinformatics 34, i884–i890 (2018).

53. Langmead, B. & Salzberg, S. L. Fast gapped-read alignment with Bowtie 2. Nat. Methods 9, 357–359 (2012).

54. Wu, F. et al. A new coronavirus associated with human respiratory disease in China. Nature 579, 265–269 (2020).

55. Pedersen, B. S. & Quinlan, A. R. Mosdepth: quick coverage calculation for genomes and exomes. Bioinforma. Oxf. Engl. 34, 867–868 (2018).

56. Li, H. et al. The Sequence Alignment/Map format and SAMtools. Bioinforma. Oxf. Engl. 25, 2078–2079 (2009).

57. Grubaugh, N. D. et al. An amplicon-based sequencing framework for accurately measuring intrahost virus diversity using PrimalSeq and iVar. Genome Biol. 20, 8 (2019).

58. Cingolani, P. et al. A program for annotating and predicting the effects of single nucleotide polymorphisms, SnpEff: SNPs in the genome of Drosophila melanogaster strain w1118; iso-2; iso-3. Fly (Austin) 6, 80–92 (2012).

59. Cingolani, P. et al. Using Drosophila melanogaster as a Model for Genotoxic Chemical Mutational Studies with a New Program, SnpSift. Front. Genet. 3, 35 (2012).

60. O‘Toole, Á. et al. Assignment of epidemiological lineages in an emerging pandemic using the pangolin tool. Virus Evol. 7, veab064 (2021).

61. Colquhoun, R. & Jackson, B. Scorpio. (2021).

62. Rambaut, A. et al. A dynamic nomenclature proposal for SARS-CoV-2 lineages to assist genomic epidemiology. Nat. Microbiol. 5, 1403–1407 (2020).

63. Jared Simpson & de Borja, R. ncov-tools. (2020).

64. Aksamentov, I., Roemer, C., Hodcroft, E. & Neher, R. Nextclade: clade assignment, mutation calling and quality control for viral genomes. J. Open Source Softw. 6, 3773 (2021).

65. Tsueng, G. et al. Outbreak.info Research Library: A standardized, searchable platform to discover and explore COVID-19 resources and data. 2022.01.20.477133 (2022) doi:10.1101/2022.01.20.477133.

66. Minh, B. Q. et al. IQ-TREE 2: New Models and Efficient Methods for Phylogenetic Inference in the Genomic Era. Mol. Biol. Evol. 37, 1530–1534 (2020).

67. Abudahab, K., Underwood, A., Taylor, B., Yeats, C. & Aanensen, D. M. Phylocanvas.gl: A WebGL-powered JavaScript library for large tree visualisation. https://osf.io/nfv6m/ (2021) doi:10.31219/osf.io/nfv6m.

68. Yu, G., Smith, D. K., Zhu, H., Guan, Y. & Lam, T. T.-Y. ggtree: an r package for visualization and annotation of phylogenetic trees with their covariates and other associated data. Methods Ecol. Evol. 8, 28–36 (2017).

69. Sagulenko, P., Puller, V. & Neher, R. A. TreeTime: Maximum-likelihood phylodynamic analysis. Virus Evol. 4, vex042 (2018).

70. Huddleston, J. et al. Augur: a bioinformatics toolkit for phylogenetic analyses of human pathogens. J. Open Source Softw. 6, (2021).

71. Seabold, S. & Perktold, J. Statsmodels: Econometric and Statistical Modeling with Python. in (2010). doi:10.25080/MAJORA-92BF1922-011.

72. Huerta-Cepas, J., Serra, F. & Bork, P. ETE 3: Reconstruction, Analysis, and Visualization of Phylogenomic Data. Mol. Biol. Evol. 33, 1635–1638 (2016).

73. Katoh, K. & Standley, D. M. MAFFT Multiple Sequence Alignment Software Version 7: Improvements in Performance and Usability. Mol. Biol. Evol. 30, 772–780 (2013).

74. Hoang, D. T., Chernomor, O., von Haeseler, A., Minh, B. Q. & Vinh, L. S. UFBoot2: Improving the Ultrafast Bootstrap Approximation. Mol. Biol. Evol. 35, 518–522 (2018).

75. BioRender. BioRender. (2022).

76. Inkscape Project. Inkscape. (2020).

77. Libin, P. J. K., Deforche, K., Abecasis, A. B. & Theys, K. VIRULIGN: fast codon-correct alignment and annotation of viral genomes. Bioinformatics 35, 1763–1765 (2019).

78. Kozlov, A. M., Darriba, D., Flouri, T., Morel, B. & Stamatakis, A. RAxML-NG: a fast, scalable and user-friendly tool for maximum likelihood phylogenetic inference. Bioinformatics 35, 4453–4455 (2019).

79. Jordan, G. E. & Piel, W. H. PhyloWidget: web-based visualizations for the tree of life. Bioinformatics 24, 1641–1642 (2008).

80. Wertheim, J. O., Murrell, B., Smith, M. D., Kosakovsky Pond, S. L. & Scheffler, K. RELAX: Detecting Relaxed Selection in a Phylogenetic Framework. Mol. Biol. Evol. 32, 820–832 (2015).

81. Hamidi, B., Wallace, K., Vasu, C. & Alekseyenko, A. V. $W_{d}^{*}$-test: robust distance-based multivariate analysis of variance. Microbiome 7, 51 (2019).

82. Alekseyenko, A. V. Multivariate Welch t-test on distances. Bioinformatics 32, 3552–3558 (2016).

83. Lung, O. et al. First whole-genome sequence of Cervid atadenovirus A outside of the United States from an Adenoviral hemorrhagic disease epizootic of black-tailed deer in Canada. 2022.02.10.479879 https://www.biorxiv.org/content/10.1101/2022.02.10.479879v1 (2022) doi:10.1101/2022.02.10.479879.

84. Fisher, M. et al. Discovery and comparative genomic analysis of elk circovirus (ElkCV), a novel circovirus species and the first reported from a cervid host. Sci. Rep. 10, 19548 (2020).

85. Kearse, M. et al. Geneious Basic: an integrated and extendable desktop software platform for the organization and analysis of sequence data. Bioinforma. Oxf. Engl. 28, 1647–1649 (2012).

86. Chatterjee, D. et al. Antigenicity of the Mu (B.1.621) and A.2.5 SARS-CoV-2 Spikes. Viruses 14, 144 (2022).

87. Chatterjee, D. et al. SARS-CoV-2 Omicron Spike recognition by plasma from individuals receiving BNT162b2 mRNA vaccination with a 16-week interval between doses. Cell Rep. 38, 110429 (2022).

88. Jennewein, M. F. et al. Isolation and characterization of cross-neutralizing coronavirus antibodies from COVID-19+ subjects. Cell Rep. 36, 109353 (2021).

89. Li, W. et al. Structural basis and mode of action for two broadly neutralizing antibodies against SARS-CoV-2 emerging variants of concern. Cell Rep. 38, 110210 (2022).

90. Enomoto, M., Bunge, M. B. & Tsoulfas, P. A multifunctional neurotrophin with reduced affinity to p75NTR enhances transplanted Schwann cell survival and axon growth after spinal cord injury. Exp. Neurol. 248, 170–182 (2013).

